# Hierarchical Optimization predicts Plasticity in the Macaque Inferior Temporal Cortex following Object Training

**DOI:** 10.1101/2024.12.27.630539

**Authors:** Lynn K. A. Sörensen, James J. DiCarlo, Kohitij Kar

## Abstract

How does the primate brain coordinate plasticity when learning to discriminate new objects? We measured consequences of object learning on macaque inferior temporal (IT) cortex, a key waypoint supporting object recognition in the ventral visual stream. Neural activity in task-trained monkeys’ IT showed increased object selectivity, enhanced linear separability across objects, and more object-invariant representations compared to task-naïve monkeys. To model these differences, we developed a computational framework using anatomically-mapped artificial neural network (ANN) models of the ventral stream with various learning algorithms. Simulations revealed that gradient-based, performance-optimizing updates of ANN internal representations accurately approximated observed IT cortex changes. These models predict novel training-induced phenomena in IT cortex, including changes independent of object identity and IT’s alignment with behavior. This convergence between empirical measurements and model predictions suggests ventral stream plasticity follows task optimization principles well-approximated by gradient descent, enabling accurate predictions about visual plasticity and generalization to test images.

## Introduction

The ability to detect and discriminate different objects is a fundamental cognitive ability that enables primates to navigate and interact with their environment effectively. However, the kinds of objects that need to be mastered may be context-dependent (e.g., identifying safe fruits to eat in a new place). As a result, learning new objects – even in adulthood – is pivotal for adapting swiftly to situational demands and thriving in unfamiliar environments. Recent years have seen much progress in computational modeling of the neural circuits that represent incoming visual images to, in turn, support the learned discrimination of objects (see (Kar & DiCarlo, 2024) for a recent overview). However, the field does not yet have computational models that explain if and how the ventral visual stream – and circuits beyond it – are modified by adult object learning.

Understanding how the brain changes to learn is a key goal in cognitive and systems neuroscience (e.g., Richards et al., 2019). However, studying the brain’s plasticity associated with a specific acquired behavior is a challenging problem that spans many spatial and temporal scales (Bredenberg & Savin, 2023; Richards & Kording, 2023). Despite much progress in developing various learning algorithms that optimize the task performance of ANNs (Guerguiev et al., 2017; Halvagal & Zenke, 2023; Lillicrap et al., 2020; Pozzi et al., 2020; Richards & Lillicrap, 2019; Rumelhart et al., 1986; Song et al., 2020) and simulations on the type of neural measurements needed for distinguishing among algorithms (Nayebi et al., 2020), the field currently lacks anatomically-precise mechanistic hypotheses and multi-level neural and behavioral measurements (Bredenberg & Savin, 2023; Lillicrap et al., 2020; Richards et al., 2019; Richards & Kording, 2023) to advance our understanding of task-specific plasticity.

We reasoned that one promising strategy for identifying plasticity is to focus on measurements in brain areas with a well-established link to behavior. The inferior temporal (IT) cortex is one such area, as it has been extensively linked to core object recognition, the ability to rapidly recognize objects at the center of gaze (DiCarlo et al., 2012). IT responses exhibit selectivity to specific types of objects (Desimone et al., 1984; Kobatake & Tanaka, 1994; Rust & Dicarlo, 2010; Yamane et al., 2008) and object-like textures (Ponce et al., 2019), are predictive of human categorization behavior (Majaj et al., 2015), and causally influence categorization behavior across a variety of tasks (Afraz et al., 2015; Kawasaki & Sheinberg, 2008; Rajalingham & DiCarlo, 2019). While IT is thus a critical sensory waypoint for rapid object categorization behavior, it remains unclear whether changes in IT are necessary to enable the learning of new objects.

Indeed, there is evidence that plasticity in IT may not be necessary for discriminating new objects. For example, various object and categorical distinctions can be retrieved from the responses of IT neurons in monkeys unfamiliar with these specific objects or with categorization tasks (Hung et al., 2005; Logothetis et al., 1995; Tsao et al., 2003). Moreover, these IT-decoded choices can fully align with human decisions on the same tasks (Majaj et al., 2015). This suggests that the behavioral change associated with object learning could be explained as a new linear readout from IT. Thus, any plasticity associated with object learning would occur downstream of IT (i.e., closer to motor outputs, Seger & Miller, 2010), in areas such as the caudate nucleus (Kim et al., 2014; Seger & Cincotta, 2005), perirhinal cortex (Pagan et al., 2013) or in prefrontal cortices (Freedman et al., 2001, 2003; Jiang et al., 2007; van der Linden et al., 2014, but see Minamimoto et al., 2010). Consequently, higher visual cortex is deemed relatively stable after reaching adulthood (Arcaro & Livingstone, 2021; Op de Beeck et al., 2008; Srihasam et al., 2012). Indeed, several studies have not observed changes in IT with category training (De Baene et al., 2008; Op de Beeck et al., 2008; Vogels & Orban, 1994). Consistent with this, a recent study modeling trial-by-trial object learning in humans (Lee & DiCarlo, 2023) demonstrated that specific learning trajectories are well-captured by dynamically updating linear decoders reading from a non-plastic IT-like feature representation of an ANN. Together, this suggests that combining versatile visual representations, as those present in IT, with plasticity in how this information is read out is sufficient for explaining how primates learn new object distinctions. From this perspective, task-specific plasticity in IT may be unnecessary or even counterproductive for task learning.

However, this view conflicts with numerous studies reporting training-induced changes in higher-level visual cortex response properties, including IT. For example, training macaques to categorize or discriminate objects can increase single-neuron selectivity for trained object discriminations relative to non-trained ones (Baker et al., 2002; Freedman et al., 2003, 2006; Kobatake et al., 1998; Logothetis et al., 1995; Pearl et al., 2023; Sigala & Logothetis, 2002), though some studies report no significant changes (De Baene et al., 2008; Op de Beeck et al., 2008; Vogels & Orban, 1994). Similarly, human neuroimaging studies have revealed increased (and faster) selectivity for trained category distinctions in the visual ventral cortex (Gillebert et al., 2009; Jiang et al., 2007; Kietzmann et al., 2016; Kuai et al., 2013; Moore et al., 2006; Op de Beeck et al., 2006; van der Linden et al., 2008, 2010). However, these effects may depend on task engagement (DeGutis & D’Esposito, 2007; van der Linden et al., 2014; Weisberg et al., 2007). Even though there is substantial variability in the reported effect sizes (Op de Beeck & Baker, 2010), and the impact of these effects on behavior is often difficult to establish, these findings raise the possibility that IT responses undergo changes to support new object discrimination behaviors, suggesting that plastic sensory representations may be necessary for object learning.

How might IT plasticity, in principle, support learning in the adult brain? Considering IT as part of a larger network of brain regions involved in object learning, rather than in isolation, it is crucial to examine how IT changes concurrently with other up- and downstream areas (Fig 1E). This perspective aligns with viewing IT as part of a distributed optimization process, where changes across the network progressively reduce a loss function to optimize performance on a novel task (e.g., (Lillicrap et al., 2020; Richards et al., 2019; Wenliang & Seitz, 2018). While there may be radically new tasks for which the entire network may need to adapt to optimize performance, in the adult brain, learning likely often recycles and slightly adapts pre-existing representations to produce new behaviors in more familiar settings. For instance, learning a new object is unlikely to induce changes in motor circuits or the retina but may necessitate adaptations in intermediate neural processing stages to ensure robust and reliable behavioral outputs. While there is extensive evidence that IT is a key node for object discrimination behavior, we do not yet know where IT lies on this continuum between stability and adaptability.

**Figure 1.**
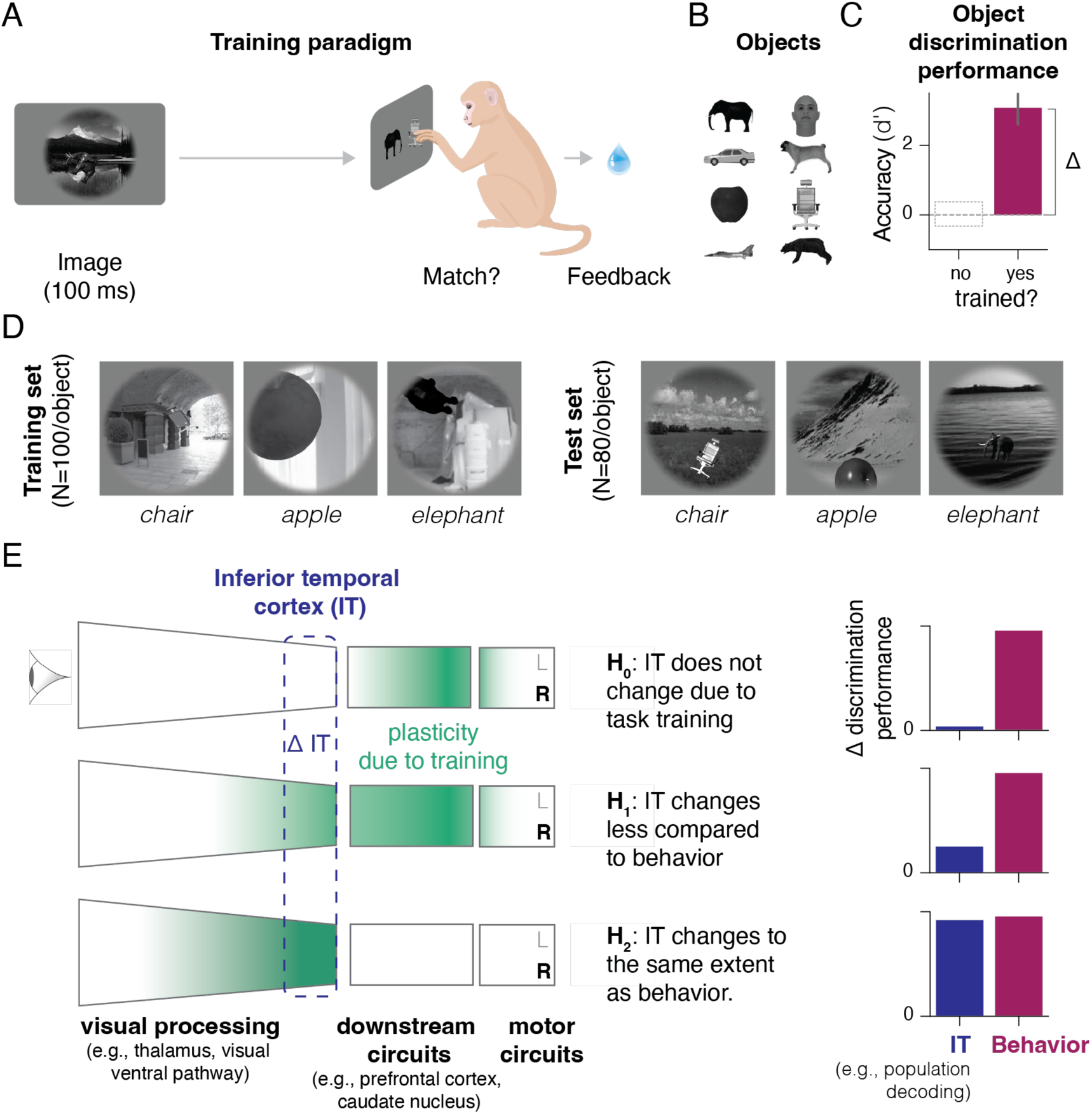
Does object training induce plasticity in sensory networks? (A) We induced task training by teaching naïve adult macaques to distinguish eight possible objects in a delayed-match-to-sample task, receiving rewards for correct choices on a trial-by-trial basis. (B) During training, monkeys learned to match object images (with different randomly generated views and backgrounds, N=800) to one of eight possible objects (elephant, person, car, dog, apple, chair, plane, bear). (C) After training, task-trained monkeys were proficient at performing this task on a new image set (N = 640). Error bars depict the 95% confidence interval of the pooled behavior of all three subjects. Δ refers to the difference between the trained and naïve group. We assume chance performance (d’ = 0) for the task-naïve group, and the dashed outline serves illustrative purposes. (D) Example images from the training and test image datasets. Test images were not presented during training and were generated using new combinations of parameters (Fig S1). (E) How are perception-action circuits adapted to accommodate such new behavioral demands? Here, we focus on a key sensory relay with a well-established link to object recognition behavior, the inferior temporal cortex (IT), and formulate three hypotheses based on prior work. One hypothesis (H_0_) is that IT does not change and that any plasticity facilitating performance on the new task happens downstream of IT (and closer to motor outputs). This view submits IT to be a downstream read-out region that no longer undergoes changes after development. Alternatively (H_1_), IT might change to support learning, yet those changes are not commensurate with those observed in behavior. Finally (H_2_), IT might reflect any change that is observable in behavior. A deeper shade of green in the drawn circuit schematic represents more plasticity to achieve the same performance level.

Focusing on IT and its link to behavior, we can discern three broad hypothesis classes (Fig 1E): One possibility is that training-induced changes occur entirely downstream of IT (such as in the caudate nucleus or prefrontal cortex). In this scenario, IT is not expected to change (*H_0_: No plasticity*) despite animals successfully mastering new objects. Accordingly, IT’s responses already afford an effective readout of the trained objects in line with the idea that IT holds task-general representations. A second possibility is that IT changes with object learning, but only to a moderate degree. According to this view, IT holds more task-related information after training in service of the observed behavioral improvements (*H_1_: Moderate plasticity*), but the most significant changes might occur outside of IT. Finally, a third possibility is that IT is the key locus for change during discrimination tasks and thus any change observed in behavior is mirrored in IT (*H_2_: Peak plasticity*). In these latter two scenarios, learning new objects would thus alter IT’s representations to a different extent when compared to behavior.

In this study, we systematically tested these hypotheses within large-scale neural recordings performed across the task-trained and naïve macaque IT cortex. Our results revealed modest but robust differences in IT representations linked to object discrimination training, supporting *H_1_*: IT recordings from animals that learned to discriminate objects demonstrated more object-selective responses at the level of individual recording sites and improved separability for the trained objects at the population level as compared to training-naïve IT recordings.

How do these differences in IT emerge, and are they indeed in service of the learned behavior? To answer these questions, we next developed an image-computable and anatomically-mapped modeling framework to test whether these training-related differences are consistent with performance-optimizing changes in a hierarchical sensory network. Using extensive ANN simulations, we demonstrate that the observed differences in IT are fully aligned with gradient-based updates to IT-like representations. Finally, we leveraged these functional models to predict two novel training-induced phenomena in the IT cortex, advancing our understanding of how IT supports learning to discriminate new objects.

## Results

As outlined above, in this study, we ask how plasticity across the macaque IT cortex enables learning of new object discrimination tasks. We aim to provide a systems-level answer to this question by examining changes in IT activity and the monkeys’ behavior in tandem with those observed in sensory-computable, mechanistic, architecturally referenced, and testable (SMART; see Kar and DiCarlo, 2024) models of the ventral stream and object recognition behavior. To achieve this, we first compare the IT activity between groups of task-naïve and task-trained monkeys across an array of single-site and population-level metrics. Next, we ask whether updating the parameters of a leading set of SMART models of the ventral stream with a set of alternative plasticity mechanisms can account for these task-training-relevant neural differences. Finally, we leverage these models to make novel predictions about IT’s reconfiguration due to task training for untrained stimulus dimensions.

### Task training enhances task-relevant information in IT

Numerous studies have investigated how plasticity in IT is linked to category and object training (for a comprehensive review, see Op de Beeck & Baker, 2010). However, while some studies suggest that training-induced plasticity increases the selectivity of IT responses (Baker et al., 2002; Freedman et al., 2003; Kobatake et al., 1998; Logothetis et al., 1995; Sigala & Logothetis, 2002), others failed to reliably detect such differences in selectivity (De Baene et al., 2008). Moreover, the magnitude of selectivity varies notably across studies (Op de Beeck & Baker, 2010), highlighting the need to revisit this question with large-scale neural recordings (Urai et al., 2022). Prior studies on IT’s plasticity have primarily focused on analyses of single-neuron properties, with limited attention to changes at the population level (but see Booth & Rolls, 1998 for an analysis of information sparseness and object familiarity and a recent study by Pearl et al., 2023). Consequently, it remains unclear how local changes in selectivity translate into increased information at the population level, which is available to downstream readout regions. To address these significant lacunae, we conducted tightly controlled experiments across two groups of monkeys (task-trained vs. naïve) and provided an in-depth analysis of how the neural representations vary between these groups as a function of task training.

We compared groups of task-naïve and task-trained monkeys (N = 3 each). Both groups learned to hold fixation to enable neural recordings. The task-trained group, in addition, was trained to discriminate objects (Fig 1B, D) using operant conditioning in a delayed-match-to-sample task (see Fig 1A for a schematic). After training, all task-trained subjects demonstrated high-performance levels on unseen images (Fig 1C). To assess the effects of task training on IT, we compared multi-unit IT responses recorded from task-naïve subjects with those of task-trained ones (Fig 2A). All recordings were performed using chronically implanted Utah arrays (2-3 arrays per monkey; see Fig S2 for array locations). During recording, monkeys passively fixated on a sequence of naturalistic images, each containing one of eight objects (Fig 2B). We presented the images in a random order, briefly (100 msec), and at the center of gaze (central 8°). To achieve comparable recording quality in both groups, we pooled IT sites (with statistically indistinguishable response reliabilities; see *Methods*) from each group (task-trained vs. naïve) by performing 1000 pseudo-random draws that ensured an equal number of recording sites per monkey (Fig 2C). Finally, we analyzed the obtained IT neuron pools for differences in task-related information using three complementary metrics: single-site object selectivity, object representational strength, and linear object decodability.

**Figure 2.**
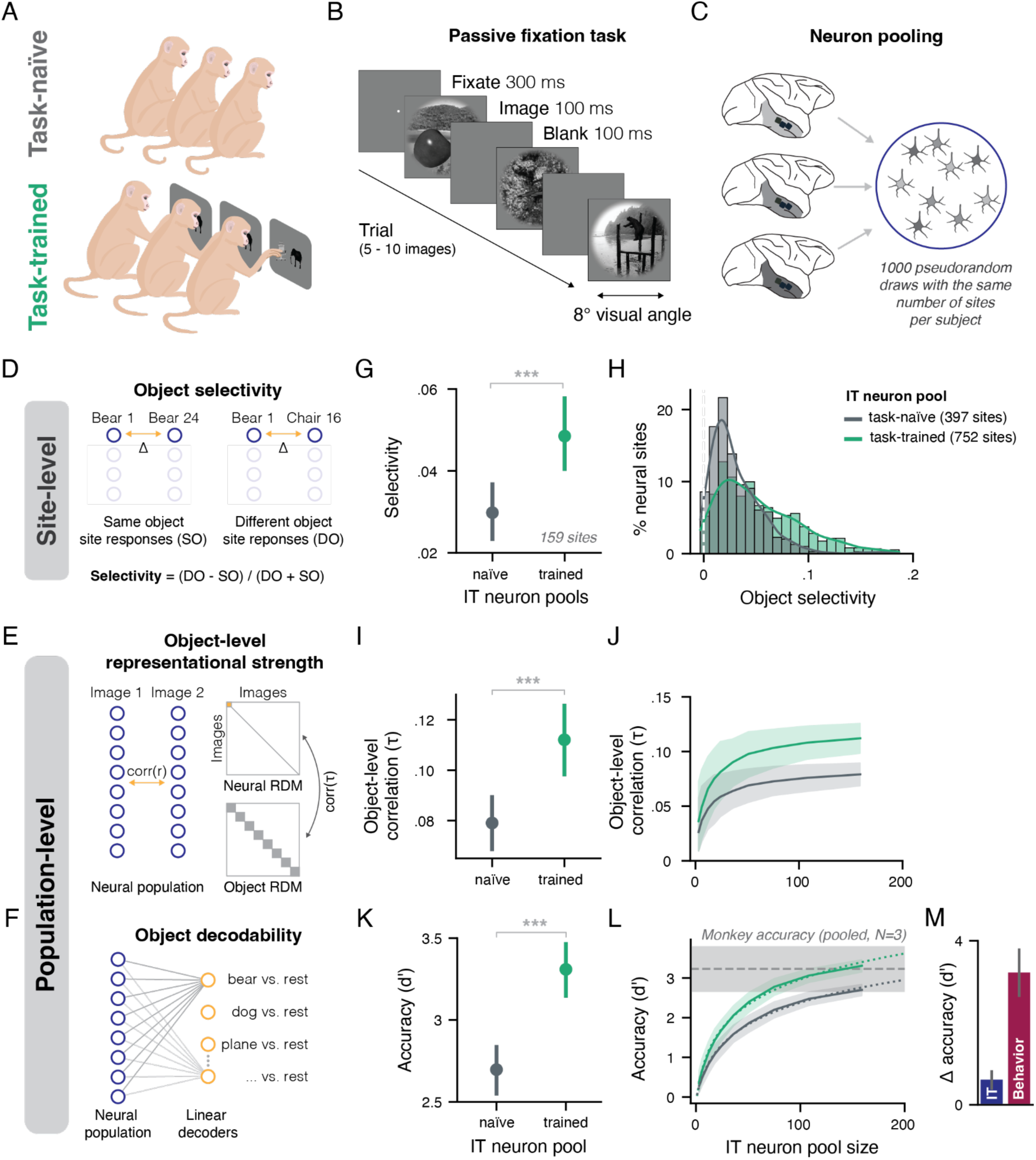
Object training enhances object-discriminating information in IT responses. (A) We recorded IT responses from groups of task-naïve and task-trained monkeys (N=3 per condition). (B) During neural recordings, all subjects passively held fixation while viewing sequences of naturalistic images. (C) We pooled all neural sites within a group using pseudo-random draws, ensuring an equal number of sites per monkey and comparable reliability across groups. (D) We assessed training-induced differences in IT responses using three complementary metrics: Single-site object selectivity captures the extent to which any site modulates its response with regard to a specific object. (E) Population-level representational strength indicates whether a neuron pool’s representational geometry is well explained by the similarity between the trained objects. We achieved this by first computing pairwise Euclidean distances across image sets to obtain the neural representational dissimilarity matrix (RDM) for a given pool. Then, we compared the neural RDM with an idealized object-level RDM to assess the representational strength using the tie-corrected Kendall’s tau distance. (F) Population-level object decodability assesses how readily the trained objects can be linearly separated in the neuron response pools. This is achieved by training one-vs.-all classifiers that find a linear hyperplane distinguishing a given object from all other objects. Compared to (E), this metric is fitted to the data (using cross-validation) and thus can be considered more powerful in assessing object-discriminating information. (G) Trained IT neuron pools (1000 draws with 159 sites each) exhibit a higher median selectivity compared to task-naïve neuron pools. The error bars depict the 95% confidence interval (CI) of the median across draws. (H) This difference is also apparent in the distribution of object selectivity across all included recording sites (without pooling). (I) Trained IT pools were more discriminating according to the trained objects in their dissimilarity structure compared to naïve pools. (J) This effect is more pronounced in larger neuron pools. (K) Trained responses hold more discriminating information about the trained objects compared to naïve responses, yielding more accurate object decodes in held-out images. (L) Left panel: The sampled IT population sizes are sufficient to approximate the behavioral performance level of the trained subjects. The horizontal shaded area shows the 95% CI of the trained monkeys’ mean accuracy (pooled, across 3 subjects) for the same images. We fit a log function to the trained IT decodes (dotted green line) to identify the crossing point of the trained decodes with the average trained monkeys’ behavioral accuracy (gray horizontal dashed line). Right panel: Comparison between differences in discrimination performance at the level of IT and behavior. IT differences show the same data as in K. All error bars and shaded areas depict the 95% confidence interval (CI) of the central tendency metric across draws.

As a first step, we compared both groups with regard to site-level response selectivity, a standard analysis for assessing training-induced neural changes (Baker et al., 2002; Freedman et al., 2003; Logothetis et al., 1995; Sigala & Logothetis, 2002). To replicate earlier findings, we computed site-specific selectivity by comparing the average pairwise response differences between images displaying the same object to those showing different objects (see *Methods* and Fig 2D for a schematic). In line with earlier studies, sites from task-trained responses showed a consistently higher degree of selectivity than those from the task-naïve pools (Fig 2G, H). This training-induced difference corresponded to a 63% increase in median selectivity (Trained: 0.048, Naïve: 0.029, CI_Difference_ (95%) = [0.008, 0.029], *p* < .001). This suggests a notable modulation of local firing rates due to task training and replicates prior claims in our larger dataset.

To address whether these site-level changes were also reflected in how task-related information is represented at the population level, we performed two complementary analyses. First, we performed representational similarity analysis (RSA, Kriegeskorte et al., 2008) to track whether task training was associated with an increased clustering according to the trained objects. Second, we fitted linear decoders to estimate the overall information about the trained objects for downstream readout.

We chose RSA for quantifying training-induced changes since it allows for quantifying changes in local geometry with regard to a specific conceptual model (i.e., the grouping of different views of the same objects). Moreover, RSA does not require any fitting procedure and thus provides an estimate of task-related information in the native data space (Fig 2E). In agreement with the selectivity analysis, we found that task-trained IT pools were more object-specific (i.e., showed lower dissimilarities within object views as compared to between different objects) than task-naïve pools (Fig 2I, Trained: 0.112 ± 0.007 (mean ± SD), Naïve: 0.079 ± 0.005, CI_Difference_ (95%) = [0.015, 0.05], *p* < .001). This difference was enhanced in larger neuron pools (Fig 2J) and corresponded to a 41% increase in the largest neuron pools (159 sites).

Finally, we asked whether these differences in local geometry translate into an overall increase in information available for downstream regions via linear read-out. We assessed this by training linear classifiers to predict the object labels (using cross-validation) based on the neural responses from both groups (Fig 2F). We observed that object predictions from the trained IT populations showed a significantly higher accuracy compared to naïve IT populations (Fig 2K, Trained: 3.31 ± 0.08 (mean ± SD), Naïve: 2.70 ± 0.08, CI_Difference_ (95%) = [0.398, 0.811], *p* < .001). Given that we could only record from a limited sample of neural sites (with respect to the total number of neurons in IT), we approximated how our estimated decoding accuracy differences scale with the number of sampled sites. As for representational strength, we observed that the object information increased in larger neuron pools (Fig 2L, 22.8% increase at 159 sites). Of note, despite the significant difference between the two groups, we observed that both IT populations could reach the behavioral performance levels shown by the trained monkeys (73 and 142 sites for the naïve and trained sites, respectively). This suggests a quantitative scaling effect of training that enhances the readout for downstream regions rather than a qualitative change due to task training (i.e., an effect through which performance only becomes possible with training). In other words, in both pools, IT holds sufficient information to guide the observed behavior, but in the trained IT recordings, the same level of performance is achieved using fewer sites. Taken together, these results complement the representational strength analysis and point to the fact that task training affected the encoding of visual information in IT at the population level, enabling a more efficient readout of task-relevant information for downstream areas.

How do these observed training-induced differences compare to differences in interindividual variation? That is, could the observed effect be driven by differences between a few subjects, irrespective of their training history? To test this, we implemented an additional analysis using a hierarchical permutation bootstrapping test. This test not only captures the variability between groups but is also sensitive to variability within both groups (see *Methods*). We observed that the effects linked to task training were significantly greater than what would be expected from interindividual variation for selectivity (*p =* .003, Fig S4A), representational strength (*p =* .041, Fig S4B), and object decoding (*p =* .002, Fig S4C). These results support that the observed differences can be attributed to task training and further help to establish that task-training-related differences can be measured in between-group designs despite interindividual variability.

Thus far, we pooled all available IT recordings within each group, irrespective of anatomical location, to maximize our signal-to-noise ratio. However, this pooling might introduce unintended differences between the task-naïve and trained groups if the recorded sites differ anatomically. Given that neural responses vary systematically along the posterior-to-anterior axis of IT (Freiwald & Tsao, 2010; Issa et al., 2018; Tsao et al., 2008) our observed group differences could result from anatomical variation rather than task training itself. To directly test for this possibility, we subdivided the recording arrays into distinct IT subregions (AIT, CIT and PIT, Fig S2) and repeated our object decoding analyses separately within each subregion. We observed that responses in all subregions contained more information on average, about object identity in the task-trained compared to the naïve group (Fig 3B). Still, the quantitative differences were only statistically different in central IT where many recording sites were available (AIT: *p* = .08,CIT: *p* < .001.001, PIT: *p* = .056). Addtionally, hemispheric differences in array locations could be another driver of training-unspecific differences in our results. Repeating our analysis for the left hemisphere only, we again observed a pronounced increase in task-related information in the task-trained IT responses (Fig 3C, *p* <.001). Lastly, we investigated whether variability in effect sizes across IT subregions was attributable to differences in the number of available recording sites. Indeed, all anatomically-constrained training effects were well predicted by effect of the number of sites in the entire dataset, suggesting that the number of sites, rather than anatomical location, was the stronger determinant of observed training effect sizes (Fig 3D).

**Figure 3.**
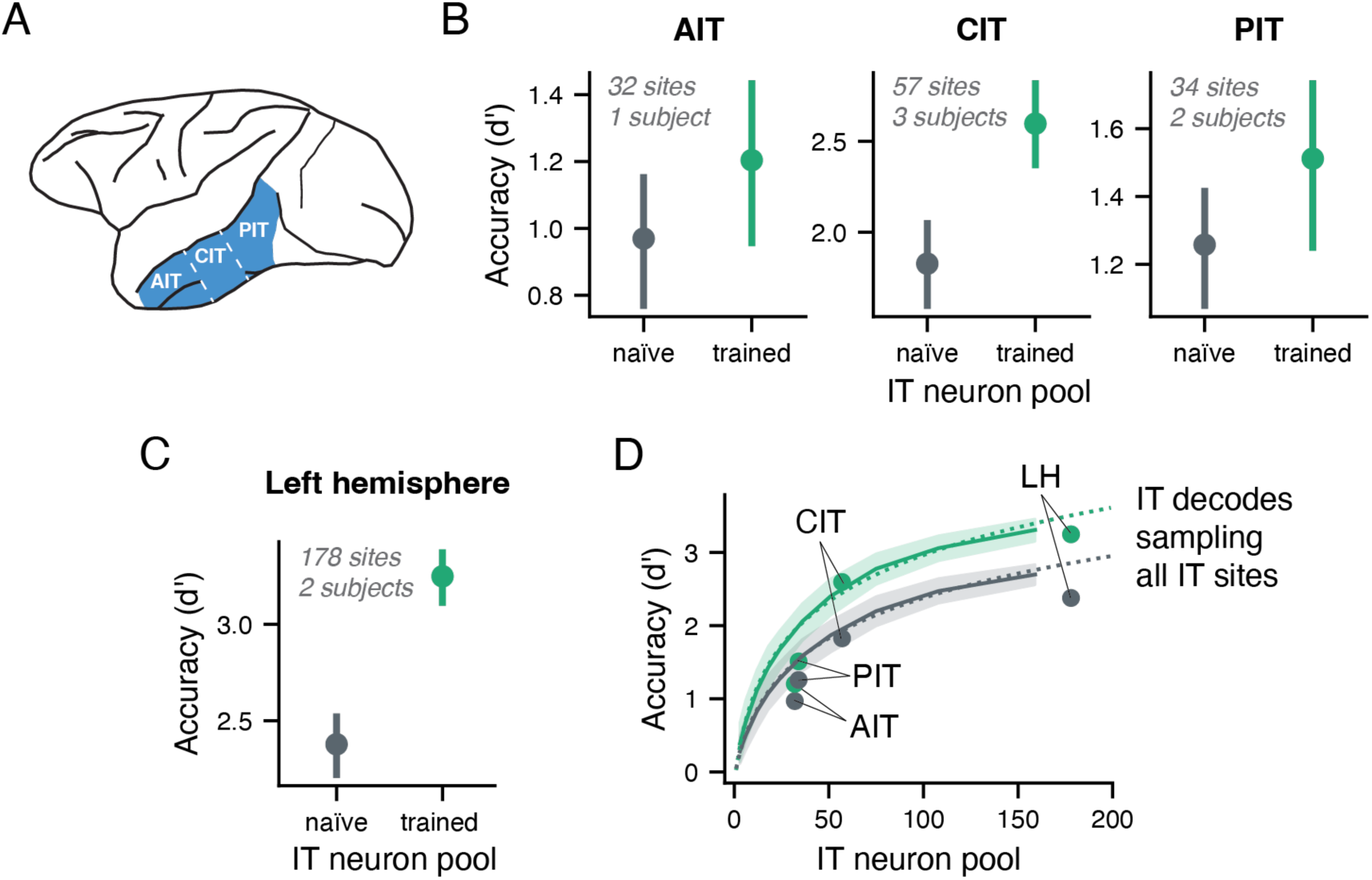
Training effects in object decoding scale with dataset set size, not anatomical location. Are the differences between trained and naïve monkey IT responses due to differences in the anatomical locations of the recording arrays? (A) To investigate this possibility, we classified the arrays from all subjects as either posterior, central or anterior IT (PIT, CIT, AIT, respectively). (B) Recordings from trained animals in all subregions of IT yield higher decoding accuracies compared to recordings from naïve subjects. As in our main analysis (cf. Fig 2K), recordings were pseudorandomly sampled to counterbalance for differences in reliability and subject counts between groups. (C) Are differences due to imbalanced sampling across the hemispheres? Repeating our analysis only in the left hemisphere, we again observe an advantage of trained IT responses. (D) The difference in effect sizes across analyses is driven by the number of available sites. Lines show decoding accuracies obtained when subsampling the entire IT dataset (irrespective of an array’s anatomical location) and the dashed line shows a fitted function to that data. Data from B and C in annotated for comparison.

In sum, task-trained IT responses displayed more object-selective site responses, more task-aligned representations at the encoding stage, and more linearly decodable information available for downstream readout. These differences are thus inconsistent with the view that IT does not change with task training (i.e., we can reject *H_0_*). How do these differences at the level of IT, compare to those observed in behavior? If training-induced behavioral differences are directly passed down from IT, we’d expect these differences to have similar magnitudes (*H_2_*). Instead, Fig 2M demonstrates that the differences at the level of IT are dwarfed by those apparent in behavior (and thus reject *H_2_*).

### Training hierarchical sensory models as hypotheses of IT plasticity

Our empirical results, as shown in the previous section, demonstrate that while task training induces task-relevant changes in IT responses, those differences appear insufficient to account for the differences in behavior (Fig 2M). These findings falsify two initial hypotheses (Fig 1E), which predicted either no changes in the IT cortex upon task learning (*H_0_*) or equivalent change to that observed in behavior (*H_2_*). If IT changes upon learning, what determines the magnitude of these differences? What are the constraints under which such moderate differences may be observed in a neural network serving the function of object recognition? How do these task-relevant differences in IT and the differences in behavior (a.k.a. learning) co-occur? To address these questions, we propose training (SMART) ANN models as an alternative hypothesis space to rigorously link differences in behavior to those observed in IT. Thus, we aim to provide quantitative predictions of the magnitude of IT’s training-induced plasticity along task-relevant metrics consistent with the observed task performance.

We assumed that a deep convolutional neural network pre-trained on a large-scale image database could serve as a proxy for a task-naïve monkey’s visual system (referred to as base models), akin to evolutionary and developmental influences providing strong priors for the adult visual system. We adopted 13 pre-trained ANNs (base models) with different model architectures and optimization objectives, including supervised, weakly supervised, and self-supervised training objectives (Fig 4A, see *Methods*). For each of these base models, we performed a two-step IT-to-model layer mapping procedure. First, we identified the model layer in all base models that best encoded task-naïve IT responses (see Fig S5 for the chosen layer and position in every base model). Second, we estimated how many ANN units in that layer corresponded to a single neural recording site (Fig S6). This gave an estimated number of units in a model’s IT-mapped layer that needed to be randomly sampled to obtain a similar level of decodable information as present in the task-naïve IT recordings.

**Figure 4.**
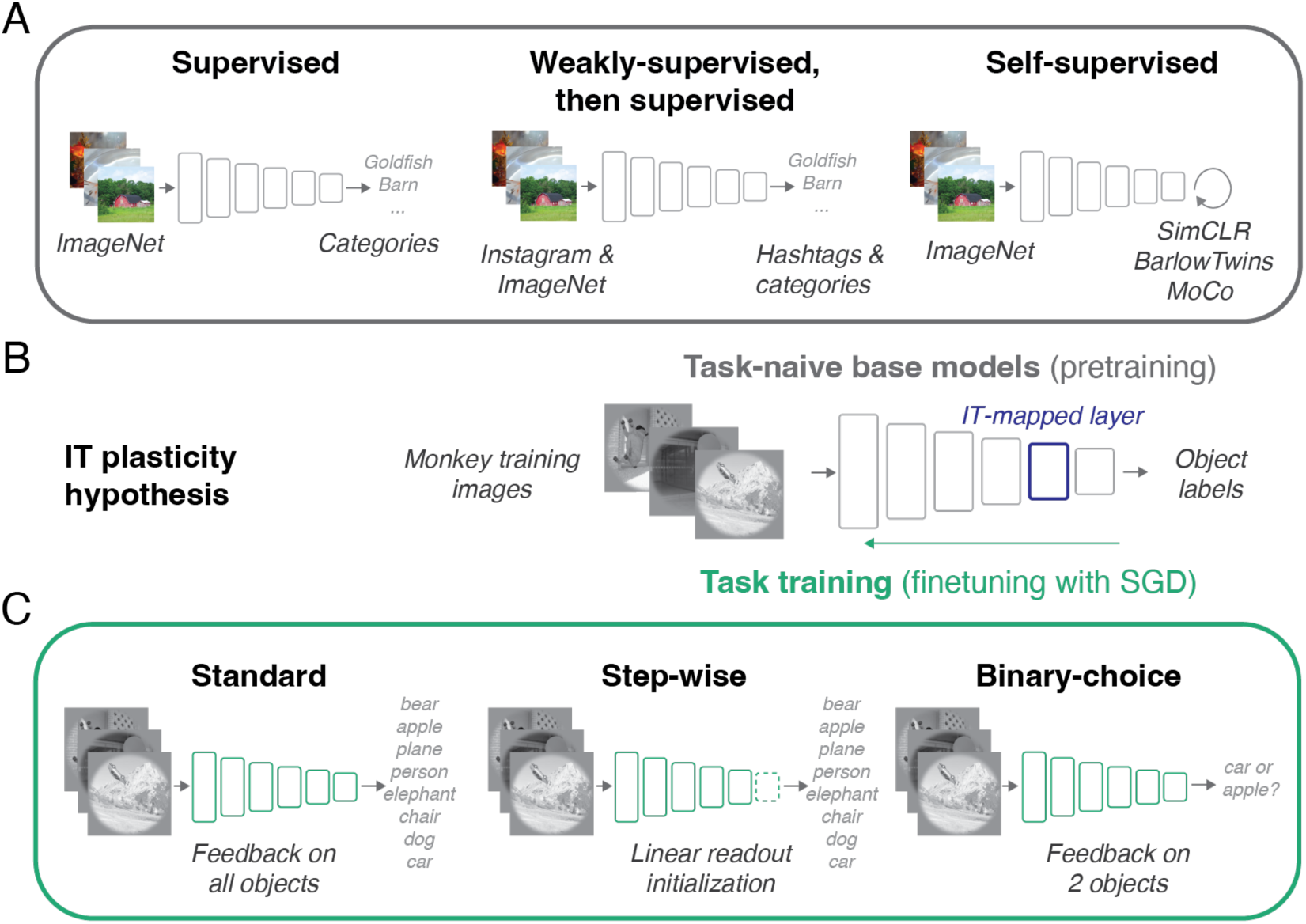
Task-optimized ANNs as IT plasticity hypotheses. We propose task optimization in ANNs as an alternative hypothesis space for IT plasticity. (A) Specifically, task-naïve IT may be readily approximated by various deep convolutional neural networks pre-trained on large-scale image databases with supervised, semi-supervised, and self-supervised objectives. We refer to these as task-naïve base models. (B) To obtain functionally relevant model hypotheses, we optimize each of these base models on the images and objects used during monkey training. Before optimization, we identify the model layer that best encodes IT responses in the task-naïve group (IT-mapped layer, Fig S5) and establish how many layer units approximate the information contained in a single IT recording site (Fig S6). (C) We apply three training strategies to every base model and repeat this procedure 20 times with various hyperparameter combinations (learning rate, regularization strength, batch size, etc.). The standard training strategy refers to replacing the final model layer with a randomly initialized layer with 8 object nodes and then fine-tuning all model weights simultaneously. For the stepwise training, we first train the linear readout layer (with the remaining network weights frozen) and then fine-tune the remaining network weights. This mimics a rapid and slow phase of plasticity. For binary-choice training, we mimic the sparse binary feedback in the monkey’s delayed match-to-sample task by only updating the model weights with regard to the target class and a randomly chosen distractor class.

With this initial mapping between task-naïve IT and base model in place, we then trained (i.e., fine-tuned) these base models using the same training images and object labels used to train the monkeys on these tasks (100 images per object, 800 images in total). To account for the possibility that certain learning strategies may be more plausible than others, we included three different training strategies: a standard fine-tuning approach, a stepwise finetuning strategy with an optimized initialization of the object readout layer, and a binary choice training that mimicked the choice and feedback structure of the delayed-match-to-sample task used during monkey training (Fig 4C, see methods for further details). During training, all model weights were free to change (i.e., no frozen parameters). In total, we trained 660 models while varying various hyperparameters (from 11 base models since 2 base models already failed during the task-naïve IT mapping stage).

### Task-optimized ANNs reproduce IT’s task-training-induced differences

As a first step, we reasoned that any eligible model of IT plasticity should perform on par with the trained monkeys (Fig 4A). Therefore, we tested all trained models for generalization on the monkey’s test images (N=640, not included in the model’s training dataset). We found that all training approaches and most base models yielded models that performed equally well as the pooled trained monkeys (Fig 4B, Fig S7A). This provided us with 208 model hypotheses and excluded under- and over-performing models from further analyses (∼32% remaining, 208/660 models). We considered these models as plausible accounts of how the observed changes in behavior could be implemented by a system learning a new task.

For each of the considered models, we then applied the same three metrics used to characterize changes in the IT recordings (i.e., change in selectivity, representational strength, and decoding, cf. Fig 2). A good model of IT plasticity should ideally match the changes in all three metrics (Fig 5A, bottom panel). On the other hand, a model that does not resemble IT along any of the metrics is likely not a good model of IT plasticity. Finally, since our evaluated metrics possibly capture different aspects of training-related changes (e.g., local vs. global changes), some models may only resemble IT in one metric but not in another. Such a pattern might reflect that each of these metrics might characterize different aspects of the neural differences.

**Figure 5.**
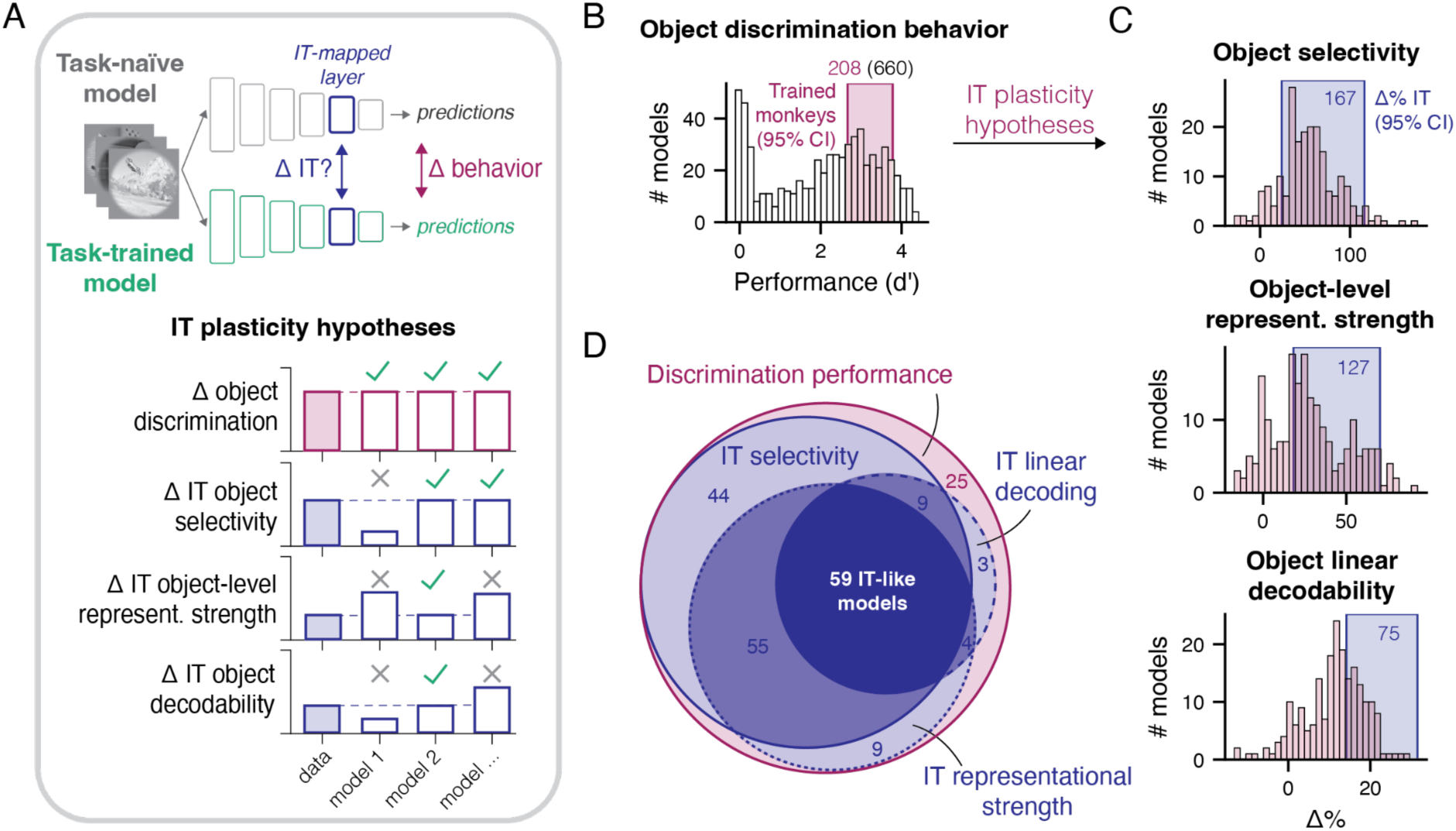
Various task-optimized ANNs reproduce IT’s task-training differences. (A, upper panel) After training, all models showed changes in their behavior, how did these changes relate to the changes in their IT-mapped layer? (A, lower panel) To create a parallel to the empirical data, we only included models that performed on par with the trained monkeys after task training (magenta-outlined bars). We then analyzed the IT-mapped layers for changes due to task training by comparing the task-trained models to their respective task-naïve base models. In particular, we compared the IT-mapped layers (task-naïve/pretrained vs. task-trained) using the same metrics as those adopted during neural data analysis (Fig 2). This step provides specific predictions of how any model’s IT-mapped layer’s activations were modified during task training to optimize its task performance. We expect a good model to reproduce the magnitude of IT changes along all three metrics (i.e., three green check marks). (B) Only a subset of task-trained models performed on par with the trained monkeys (magenta bands show the 95% CI of the pooled monkey performance on held-out test images (N=640). Training-specific results are shown in Fig S7A. We only evaluated models with matching test performance as relevant models of IT plasticity. (C) For all metrics, we saw a significant proportion of models displaying change magnitudes akin to those in IT recordings. The x-axis shows the mean estimate of changes across unit draws for every model (N = 208). Most, but not all models showed an increase in these object-related metrics due to task training (i.e., distribution is shifted right of 0). Blue shaded areas depict the 95% confidence interval of observed changes in IT (across 1000 draws). The annotated number describes how many models matched the neural data in their magnitude and direction of change for a given metric. (D) Agreement across metrics. For every model matching IT along a single metric, we ask whether the same model matched the neural data for the other two metrics. By design, all models matched the object discrimination performance (magenta circle). The darkest shade of blue shows the proportion of models matching all metrics. The medium and lighter shades of blue depict the proportion of models matching two or only a single metric.

For all metrics, a substantial proportion of tested models showed the same level of changes upon object learning in their IT-mapped layers as observed in IT (Fig 5C, 80%, 61%, and 36% for object selectivity, representational strength, and decodability, respectively). Inspecting the overall model distribution suggests a general alignment in terms of magnitude with IT changes. This alignment was particularly pronounced for changes in selectivity. This suggests that training ANNs on the same task mastered by the task-trained monkeys and observing the comparable levels of task proficiency, yields, in most cases, models that mimic the task-related differences observed in IT cortices.

Were there any models that exhibited IT-like changes across all metrics? Among all 208 assessed models, we identified 59 models that were indistinguishable from the neural data for all tested metrics (referred to henceforth as “IT-like” models). Moreover, Fig 5D highlights how one metric can be considered more sensitive to identifying IT-like models. For example, observing a match in object decoding appeared to be a good predictor for a match in the remaining two metrics. On the other hand, many models resembled IT’s changes in selectivity, yet these models did not necessarily also match IT’s changes in population-level metrics.

Can we identify factors that explain why one model better accounted for the changes in IT than another? To investigate this, we returned to the variations introduced during task training (i.e., base models and task training strategies, Fig 4). We assessed whether any training strategy or base model was associated with a higher proportion of IT-like models (i.e., a match in all three metrics). Despite the differences in how the networks were fine-tuned, we observed that all three training strategies produced IT-like models in a relatively comparable proportion (Fig S7B), thus suggesting that the larger changes associated with task optimization using gradient descent led to the IT-likeness. In the tested base models, we further observed that ResNet-style architectures were more effective at producing IT-like models compared to VGG-style and AlexNet architectures, pointing to a potential benefit of skip connections for producing IT-like models. We considered base models with different pre-training objectives, reasoning that models pre-trained using a self- or weakly-supervised objective could be a more appropriate starting point for IT-like task training. In particular, adult naïve monkeys will have received very little explicit (i.e., supervised) feedback about object categories (Zhuang et al., 2021). Thus, adopting a base model exposed to a similar feedback regime as the task-naïve monkeys could be critical for explaining the observed differences in IT. Against our expectation, we observed that self-supervised and supervised pre-trained models produced relatively comparable proportions of IT-like models (except ResNet50 pre-trained with MoCov2). For the two tested weakly-supervised models, we could not identify a unit-mapping scaling that approximated naïve IT recording sites. These results indicate that the exposure to category signals during pre-training does not result in less IT-like models.

### IT-like models best predict differences along untrained dimensions in IT

Beyond a fit to the training-related differences, a good model of IT plasticity should also generalize and predict novel phenomena beyond the scope of training. In the previous section, we identified models fully mimicking the effects of object discrimination training on both behavior and IT responses. Yet, it remains possible that such IT-like models exclusively mimic object-specific changes (because they were selected for it, cf. Fig 5) but fail to predict anything beyond the direct effects of training. A way to further assay the quality of our models is thus to probe whether IT-like models generalize to other domains or dimensions that were not reinforced during training. Examples of such domains are predicting an object’s position, eccentricity, rotation, and an image’s perceptual difficulty reflected in the trained monkeys’ behavior. Examining decoding differences along such dimensions is meaningful since the effects of object discrimination training (e.g., a larger separation between images belonging to separate objects, Fig 6A, upper panel), are likely also to affect other stimulus-related dimensions (e.g., an object’s size, Fig 6A, lower panel). Such untrained dimensions can thus serve as additional probes for examining if two systems underwent a functionally similar change (from the perspective of downstream readout regions from IT). By leveraging a variety of these training-unrelated decodes, we can thus put our identified IT-like models to the critical test: If such models do not only predict IT’s object-related differences but also generalize to a variety of other untrained decodes, this provides strong evidence that these models capture more general aspects of training-induced IT’s change (Generalization, Fig 6B). Alternatively, suppose a model’s object-specific IT-likeness shows no correlation or even an inverse relationship with its ability to decode untrained image aspects. In that case, this suggests that the model captures only narrow, object-specific differences. This latter case could be considered a form of overfitting to the empirical benchmarks tested thus far (No link or overfitting, Fig 6B). In what follows, we tested the ability of IT-like models to generalize to these untrained readout dimensions. We also evaluated less IT-like models as control conditions to understand whether a model’s generalization directly stems from its IT-likeness.

**Figure 6.**
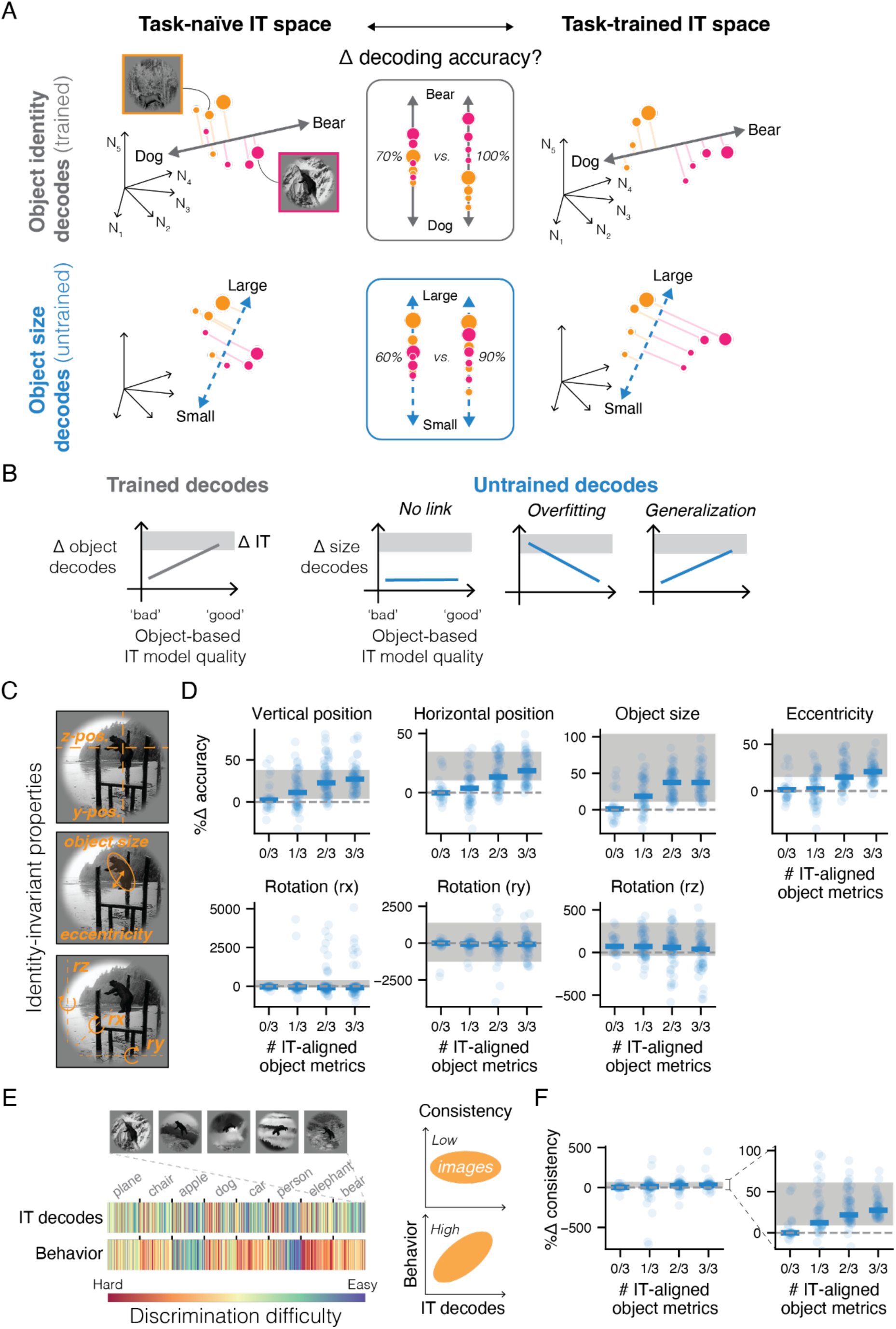
IT-like models generalize to changes along untrained readout dimensions. (A) We investigate how object discrimination training affects IT representations by comparing linear decodable information in task-naïve versus task-trained IT responses. We visualize this using 8 example images in high-dimensional IT space, with each dimension representing a neuron’s response. Orange circles represent dogs and pink circles represent bears, with circle size indicating object size. Bidirectional arrows show readout dimensions for different decoding tasks. The upper panels show how object identitydecoding improves after training, with increased separation between object classes (along the readout dimension). The lower panel applies an untrained object size decoder. A linear decoder’s accuracy will depend on how the images are encoded. Differences in encoding due to task training can thus be quantified using untrained decoders as probes. (B) By design, models resembling IT in their object identity-related metrics (‘good models’) will best capture the observed differences in object identity decoding, whereas ‘bad’ models will not predict such differences (left). Can such good models also best predict other untrained dimensions? We hypothesize three scenarios of how a model’s (object-related) IT-likeness relates to its predictivity for untrained changes: (1) There may not be a link, indicating that a model’s fit based on object metrics does not transfer to other dimensions. (2) There could be a negative link. This means that the ‘best’ models are the worst for predicting differences in other domains. This pattern of result is indicative of overfitting on the object identity-based differences. (3) There could be a positive link suggesting generalization between a model’s ability to predict object identity-related differences to other untrained differences. (C) Beyond an object’s target class, test images also varied along identity-invariant properties such as an object’s location (horizontal and vertical position, eccentricity), size, and pose (x-, y-, and z-axis rotation). (D) Most IT-like models best predict these identity-invariant changes. We evaluated four groups of models with increasing IT-likeness as assessed via a match in object identity-related change metrics. The x-axis spans models with no indication of IT similarity to those matching IT for all three object-related change metrics (cf. Fig 5D). Shaded gray bars show the 95% CI of the observed changes in the neural data. Individual dots show each model within a group. Blue horizontal bars depict the median change across all models. (E) To what extent do training-related IT changes increase the alignment with image-specific (trained) behavior? To obtain image-specific performance estimates for the IT recordings, we fit linear decoders to the neural recordings and estimate image-level decoding accuracy (d’, see Methods) for a specific image using held-out predictions. Intuitively, image-level decoding accuracy captures the difficulty in correctly assigning the target object to a given image while accounting for the false alarm rate associated with that target object. We can estimate the same metric based on the monkey’s choices during a behavioral experiment. (F) Most IT-like models best predict the empirical increases in IT-behavior consistency resulting from task training. The right panel is a zoomed-in version of the left panel.

A prior study (Hong et al., 2016) demonstrated that object-orthogonal properties, including an object’s location, size, position, and rotation from a canonical view (Fig 6C), can be linearly decoded from task-naïve IT responses. The accuracy of these IT-based decodes was, in turn, correlated with that of human judgments, suggesting that IT may serve as a readout region for such judgments. This makes these dimensions ideal candidates for assessing generalization in the IT-like models. In particular, we expect that models with the closest alignment in object-related IT metrics will also most accurately predict the training-induced IT differences for each of these object-invariant dimensions. To test this, we leveraged that our experimental images were created by rendering a 3D object with varying poses, locations, and sizes (Fig S1). Consequently, we had access to the ground-truth values of these image properties and could thus quantify how accurately they could be decoded from IT responses. Concretely, we fit cross-validated linear decoders to the pooled responses in both groups (similar to object decoding) to measure linear decoding accuracy (see *Methods*).

For the IT recordings, we found that object training significantly increased identity-invariant information about an object’s position, size, and eccentricity (all *p* < .005) but not about its rotational attributes (all *p >* .131, Fig S8A). Good models of IT plasticity were thus expected to mimic differences for some dimensions (object’s position, size, and eccentricity), yet not for others (rotational properties). Then, we performed the same analysis in each of the models’ IT-mapped layers. We evaluated both IT-like models (i.e., matching all three object-related metrics) as well as less IT-like models with either two, one, or no match with any of the object-related metrics (Fig 5D) to isolate the effect of IT-likeness. For the properties that significantly differed between task-naïve vs. -trained IT recordings (i.e., object location, size, and eccentricity), we observed that IT-likeness introduced significant differences between model groups (upper panel, Fig 6D, *H(3) =* 38.34 - 51.03, all *p* < .0000001) with IT-like models always producing a larger increase in accuracy compared to IT-unlike models. Moreover, an increasing degree of IT likeness was associated with a better match with the empirically observed differences. For properties that did not differ between the task-trained vs. task-naïve neural recordings (i.e., rotations), we observed that all model groups provided good approximations of the null effect (lower panel, Fig 6D). We also did not observe a significant difference between IT-like and IT-unlike models for these properties (*H*(3) = 0.32 - 7.51, all *p* < .057). This pattern of results supports the notion that IT-like models capture more general training-induced differences that extend beyond object-related metrics.

In addition to these object identity-invariant properties, a prior study demonstrated that an image’s behavioral difficulty is readily decodable from IT responses (Majaj et al., 2015). That is, the degree to which an image’s object is linearly distinguishable from IT responses is linked to the behavioral difficulty of picking the correct target canonical view when presented alongside various distractor objects (Fig 6E). While we have shown in our prior analysis that there is an overall increase in mean decoding accuracy (Fig 2K), we do not yet know how the link between image-by-image decoding patterns differs with task training and how these IT-decoded image-by-image patterns, in turn, compare to behavioral patterns. Importantly, we can adopt this behavioral metric as another readout dimension to further probe the IT-like models for generalization. As with the object identity-invariant properties, we again expect that IT-like models should most accurately predict any effects of task training, whereas less IT-like models are not expected to predict these differences.

In the IT recordings, we observed a significant increase in behavioral consistency between task-trained and task-naïve IT recordings (Trained: 0.56 ± 0.03 (mean ± SD), Naïve: 0.43 ± 0.03, CI_Difference_ (95%) = [9.86, 60.18], *p* = .001, Fig S8B). Thus, image-level behavior was better predicted using task-trained compared to task-naïve IT recordings. Next, we tested how models with varying levels of IT-likeness can predict these differences in behavioral consistency. We found that there were indeed significantly different predictions according to a model’s IT-likeness (right panel, Fig 5F, *H*(3) = 29.24, *p* < .00001), with IT-like models predicting the most accurate increases in IT-behavior consistency and IT-unlike models predicting no increase. In line with the object identity-invariant properties, this thus provides additional evidence that the evaluated IT-like models capture more general differences that extend beyond changes in the encoding of object-related information, reflecting both encoded image latents as well as fine-grained behavioral characteristics.

While IT-likeness was the strongest factor shaping these results, we also noticed that all IT-likeness groups featured several outlier models (Fig 6F, left panel). An exploratory analysis revealed that such extreme values were exclusively linked to models trained using binary choice feedback but not using the other two training approaches. Thus, while the IT-mapped layer changes did not distinguish between training strategies (cf. Fig 5), at the behavioral level, binary-choice training produced qualitatively different image-by-image behavior despite comparable mean performance levels. A follow-up analysis further indicated that both standard and step-wise training produced highly similar image-by-image behavioral patterns. These two groups were also highly similar to the trained monkeys. In contrast, binary choice-trained models formed a distinct cluster (Fig S9). On the whole, this means that against our expectation, binary-choice training did not produce more monkey-like behavioral patterns, and instead, the other training strategies were better suited for obtaining models that resemble monkey behavior, on average, and at an image-by-image grain.

## Discussion

How does plasticity in the adult brain enable the learning of new objects? We here provided a systems-level answer to this question by examining differences in IT activity and the monkeys’ behavior in tandem with those observed in sensory-computable, mechanistic, architecturally referenced, and testable (SMART; see Kar & DiCarlo, 2024) models of the ventral stream and object recognition behavior. Specifically, we quantified the differences evoked by object discrimination training by comparing the IT responses of training-naïve monkeys with those of monkeys trained to discriminate objects. We identified modest, yet robust differences, at the local and population level, induced by object discrimination training (Fig 2, 3): IT responses from task-trained monkeys were more object-selective at the single-site level. At the population level, IT’s responses contained overall more information about the trained objects. How are these changes in IT connected to the behavioral improvements after training? To answer this question, we next combined SMART models with task optimization via gradient descent to model the monkeys’ training and to study the effect of task training on IT responses (Fig 4). Extensive model simulations revealed that the observed training-induced differences in IT were fully consistent with those of a hierarchical sensory network accommodating a new object discrimination behavior (Fig 5). Moreover, the direction and magnitude of the empirically observed IT differences could be modeled using various architectures, pre-training objectives, and training strategies. To further assess the generality of our framework, we next probed whether models closely mimicking IT (i.e., IT-like models) could predict novel training-related phenomena. Indeed, we found that IT-like models predicted the specific increases in information in IT responses about object identity-invariant properties (Fig 6). Moreover, IT-like models accurately captured the increases in IT-behavior alignment at the image-by-image grain (Fig 6). Collectively, these results support the idea that training-induced IT differences are a direct consequence of IT’s functional role and position within the visual ventral stream.

### Adopting a systems-level perspective clarifies IT’s role in object learning

A key insight from our study is that object discrimination training changed IT responses to a moderate extent yet the magnitude of these differences did not fully account for the behavioral improvements observed after training (Fig 2M). This pattern of results rejected two initial hypotheses (Fig 1E): (1) That IT does not change at all, since IT’s responses already held sufficient visual representations to support performance on the trained task (*H_0_*) and (2) that IT responses serve as a direct readout for behavior and, in turn, IT would reflect any changes apparent in overt behavior (*H_2_*). Contextualizing IT’s difference with those in behavior, enabled us to show that neither of these scenarios applied: IT’s responses were enriched in favor of the trained objects, while at the same time, not differing to the same extent as behavior. This suggests that most plasticity is expected to occur downstream of IT.

This raises the question of why IT changed to the degree that we observed and why not more or less. To investigate this, we proposed a computational framework to simulate the plasticity needed at the level of the IT cortex to support the task performance shown by the trained monkeys. These simulations revealed that IT’s change magnitude across various metrics naturally followed from its functional role within the visual hierarchy. In other words, viewing IT as part of a larger network that gradually changes in a distributed fashion to optimize task performance, reproduces the direction and magnitude of differences observed in IT responses. This perspective submits that IT’s plasticity will depend on the training stimuli, the task demands (i.e., the similarity between objects), IT’s functional state before training, and a subject’s task performance after training. This view roughly aligns with proposals of perceptual learning in IT as part of a hierarchy of representations (Riesenhuber & Poggio, 1999; Sheinberg & Logothetis, 2002; Wenliang & Seitz, 2018), but critically we here turn this proposal into a testable hypothesis grounded in image-computable models and task performance. Using our simulations, we reason that differences in the visual features of the training stimuli, their linear decodability in IT before training, and an animal’s trained performance after training may have contributed to the diversity of effects sizes reported in prior studies (see Op de Beeck & Baker, 2010)for a review), with some studies reporting sizable changes with category training (Baker et al., 2002; Freedman et al., 2003, 2006; Kobatake et al., 1998; Logothetis et al., 1995; Pearl et al., 2023; Sakai & Miyashita, 1994; Sigala & Logothetis, 2002) and others finding small or absent changes (De Baene et al., 2008; Op de Beeck et al., 2007, 2008; Vogels & Orban, 1994). Taking these factors into account when comparing across studies along with generating training-specific computational hypotheses to guide experiments has the potential to provide further insights into the circumstances under which IT may or may not change with task training.

At first glance, our neural findings seem at odds with studies reporting that human discimination behavior is already fully accounted for by task-näive IT decodes (Majaj et al., 2015) and that behavioral learning is well accounted for by leveraging general-purpose visual representations (Lee & DiCarlo, 2023). Interestingly, our analyses revealed an increase in information efficiency in task-trained compared to task-naïve IT responses (Fig 2L). This clarifies that task-trained responses do not fundamentally differ from task-naïve ones, yet subtle differences in IT associated with task training might enable a more efficient downstream readout. It is thus possible to account for discrimination behavior using task-naïve IT recordings, yet fewer downstream projections are necessary to achieve comparable results in task-trained IT responses. This highlights the role of anatomical considerations in future studies.

### Beyond IT: The Role of Downstream Networks in Object Learning

How does behavioral change emerge during object training, enabling primates to go from chance-level performance to higher accuracies? The modest changes we observed in IT, combined with evidence that object-relevant information is already present in naïve IT, suggest that learning may primarily involve optimizing how IT representations are mapped to task-relevant decisions. In our delayed-match-to-sample task, this mapping requires several computations: maintaining the sample image in working memory, comparing it with test stimuli, and translating the comparison into a motor response (left/right choice). These computations likely engage multiple brain regions beyond IT. At present, we lack task-computable SMART models for these downstream subnetworks. Notably, the IT-mapped layers in the evaluated base models were typically in the last third of the model (Fig S5) and not at the final layer. An exciting avenue for follow-up research could be to expand current models beyond the visual ventral stream. Indeed, previous work provides important clues about these downstream networks. The prefrontal cortex, particularly vlPFC, plays a crucial role in rule learning and working memory (Freedman et al., 2001; Mansouri et al., 2020; Rainer & Miller, 2000). Moreover, direct manipulation of vlPFC activity causally impacts discrimination behavior and modulates IT responses (Kar & DiCarlo, 2021). The perirhinal cortex may help transform IT object representations into more categorical formats (Pagan et al., 2013). Additionally, structures like the caudate nucleus show learning-related changes during category training (Seger & Miller, 2010). A promising direction for future research is to test whether the magnitudes of neural changes induced by object vs. category training exceeds what would be expected based on the visual dissimilarities between objects. Such an analysis would clarify the extent to which our proposed framework generalizes to and captures plasticity induced across diverse task structures. In line with prior findings (e.g., Freedman et al., 2003), our modeling work also predicts that these downstream areas of IT should demonstrate higher degrees of plasticity (as compared to IT) due to their closer connection to generating trained task behavior. How distributed, task-specific, and engagement-dependent such downstream systems are in the brain is an important avenue for future research. Extending our SMART modeling framework (Kar & DiCarlo, 2024) to include downstream networks could help bridge this gap, generating concrete hypotheses about how different brain regions coordinate plasticity during learning.

### Understanding What Makes Models Match IT Plasticity

Our analysis revealed that approximately 30% of all performance-matched models – that could perform the behavioral tasks with the same accuracy as trained monkeys – fully reproduced the training-induced differences observed in the macaque IT responses. This finding raises a crucial question: what determines whether a model will capture the hallmarks of object training in IT? Contrary to our initial hypotheses, we found that no single factor could reliably predict a model’s ability to match IT changes. While certain architectures and training strategies showed higher success rates (Fig S7), the relationship between model design and IT-like plasticity proved more complex than anticipated. For example, we included self-supervised pre-trained models in our set of base models, reasoning that a model’s prior exposure to categorical feedback during its pre-training may be an important determinant of how such a system responds to the introduction of object feedback (Zhuang et al., 2021). However, we found no systematic advantage of self-supervised over supervised pre-training (Fig S7). These results challenge the assumption that specific, biologically motivated design choices will automatically improve a model’s alignment with the brain (or its changes with training). Instead, they highlight that IT-like plasticity emerges from complex interactions between a model’s initial state, architecture, and learning dynamics.

If modelers’ intuitions about the right design choice offer no direct guidance, how can we progress in developing useful computational models of the brain to motivate future experiments? Our findings underscore the critical importance of empirical benchmarks in this computational modeling effort (Schrimpf et al., 2020). By requiring models to simultaneously match behavioral performance and neural differences we established stringent criteria for evaluating different computational hypotheses. This approach revealed that IT-like object learning systems can arise from various combinations of factors, with successful models sharing similar functional properties despite architectural differences. The predictive power of these models extended beyond our initial benchmarks to novel phenomena (Fig 6), validating their capture of fundamental principles underlying IT plasticity. However, important questions remain about the apparent diversity of successful models. Do they truly implement equivalent solutions to the object learning problem, or are our current benchmarks insufficient to distinguish between different computational strategies? While, to our knowledge, this study is among the largest investigations of such effects in non-human primates, future work with larger samples and more targeted model-separation-driven measurements may help discriminate between competing model classes. Such research could further constrain the space of viable computational theories and potentially reveal previously unrecognized principles of neural plasticity.

### Gradient Descent as a Window into Neural Plasticity

Our success in modeling IT plasticity using gradient descent raises important questions about the relationship between artificial and biological learning. While gradient descent may not mirror biological learning mechanisms, our results suggest it captures fundamental principles about how hierarchical networks optimize behavior. Our network simulations suggest that a model of IT’s plasticity is readily obtained by combining SMART models with task optimization via gradient descent to mimic object training. We here viewed gradient descent as an effective tool to simulate a normative perspective on the observed neural changes. In other words, if one wants to understand how a distributed system *should be* adapted in service of task performance, gradient descent provides the optimal solution to achieve this (Bredenberg & Savin, 2023; Richards & Kording, 2023). From this perspective, the success of our modeling approach may not be surprising. Viewing our results through this lens, we found that IT plasticity appears to follow principles of efficient task optimization, where changes in sensory representations cannot be understood in isolation from behavioral demands. The magnitude and distribution of observed plasticity reflect both computational constraints and network architecture. However, the relationship between gradient-based learning in artificial networks and plasticity in biological circuits requires careful consideration. There remains a possibility that the changes achieved via gradient descent, even though they are optimal, are not achievable in biological brains. While gradient descent computes optimal weight updates using non-local information propagated backward through the network, biological circuits face more severe constraints (Lillicrap et al., 2020; Song et al., 2020). Neural plasticity likely operates with more constrained mechanisms than artificial networks, such as predominantly local information at synapses (Sacramento et al., 2018), temporally sparse and delayed feedback signals (Guerguiev et al., 2017), and circuit motifs that may restrict information flow (Lillicrap et al., 2020; Richards & Lillicrap, 2019). For example, the structure of neural circuits could limit how task-optimizing error updates may be approximated and propagated throughout the system affecting the distribution and magnitude of changes in IT. Remarkably, our results suggest these biological constraints may not critically impact the pattern of plasticity at the scale of IT responses and behavior. We found that multiple learning algorithms, operating under different constraints, could produce changes matching empirical observations. This implies that the observed IT plasticity may reflect general principles of hierarchical learning rather than specific implementation details. Nevertheless, it is conceivable that these factors will become important when modeling inter-areal coordination, learning dynamics over time, individual differences in learning trajectories, and the transfer of learning to novel tasks. Future work combining our normative modeling approach with more biologically detailed implementations may help bridge the gap between artificial and biological learning mechanisms.

### Limitations and Future Directions

While our study provides insights into how object learning shapes IT responses, several important limitations suggest promising directions for future research. A central limitation is our reliance on between-subject comparisons rather than tracking learning-induced changes within individual subjects. Though a within-subject design would be ideal for understanding how category learning modifies neural circuits, two key challenges justified our between-subject approach. First, characterizing learning-induced plasticity within individuals requires precise measurement of baseline states and careful separation of true plasticity from measurement variability. Current techniques for longitudinal neural recordings face significant challenges in maintaining stable measurements across extended periods, making it difficult to definitively attribute response changes to learning rather than recording instability or sampling differences (Chestek et al., 2011; Dickey et al., 2009). While recent advances in chronic recording methods and statistical frameworks for analyzing individual variability (Oby et al., 2019) show promise, the field still lacks robust approaches for tracking fine-grained neural changes over learning timescales. Second, our between-subject design enabled the collection of a larger dataset (six subjects) with better statistical power than would be possible in a within-subject study given current technological constraints. This approach revealed consistent training-induced changes across subjects, providing strong evidence for systematic plasticity in IT during category learning shared across individuals. Nevertheless, future studies combining improved longitudinal recording methods with our computational framework could reveal important insights about individual learning trajectories that our between-subject design could not address.

Beyond tracking individual changes, understanding object learning will require examining how multiple brain regions coordinate plasticity. A key strength of our approach was leveraging the well-established quantitative link between IT responses and object recognition behavior (Kar et al., 2019; Majaj et al., 2015). This link provided a crucial advantage: we could systematically evaluate plasticity by measuring how well neural changes predicted behavioral improvements. Unlike many studies of neural plasticity where the relationship between neural changes and behavior remains unclear, our understanding of IT’s role in visual recognition allowed us to test specific hypotheses about how learning modifies neural circuits in service of behavior. Future work should leverage similar brain-behavior relationships to examine how regions like the prefrontal cortex (Kar & DiCarlo, 2021) and perirhinal cortex (Pagan et al., 2013) interact with IT during different kinds of learning (e.g., at the level of objects and categories). Our modeling framework could help generate specific predictions about plasticity in these regions, guiding experimental design and data analysis.

Another crucial direction is understanding how learning unfolds across different timescales and its interplay with unsupervised learning (Jia et al., 2021; Li & DiCarlo, 2008, 2010; Li & Dicarlo, 2012; Zhong et al., 2024) and pre-existing perceptual expertise (Collins & Behrmann, 2020; Martens et al., 2018; Tanaka & Taylor, 1991). Our study examined the end state of object training, but important questions remain about how representations evolve during learning. How do unsupervised learning and existing perceptual expertise facilitate the observed difference in IT responses? How do early and late stages of learning differ? What is the interplay with other nearby structures such as the hippocampus in different stages of training? Answering these questions will require new experimental paradigms that can track neural changes at multiple temporal scales while maintaining measurement stability as well as computational models that can express these different hypotheses.

## Methods

### Subjects

The non-human subjects in this study were 6 male rhesus monkeys (Macaca mulatta; 5-10 years of age). We grouped these monkeys as trained (monkeys M, N, and B) and naïve (monkeys C, T, and S) with respect to a set of object categorization tasks. We could perform a rigorous comparison across such a large cohort since all included monkeys also participated in other studies (data and methods from Monkey C and T have been described in (Majaj et al., 2015); those from Monkey N, M, and S in (Bashivan et al., 2019); Monkey N and B in (Kar & DiCarlo, 2021); and Monkey M and N in (Kar et al., 2019). All surgical and animal procedures were performed in accordance with National Institutes of Health guidelines and the Massachusetts Institute of Technology Committee on Animal Care.

### Naturalistic object stimuli

We generated synthetic (“naturalistic”) object images using free ray-tracing software (http://www.povray.org, see Fig 1D for examples. Images contained 8 possible objects (bear, elephant, face, apple, car, dog, chair, plane, Fig 1B). Each image was rendered from a 3D object model (purchased from TurboSquid) into a 2D projection while varying an object’s position (horizontal and vertical position), rotation (in the horizontal, vertical, or depth plane), and size (Fig S1A). We then added the rendered object view to a randomly chosen natural image background (indoor and outdoor scenes obtained from Dosch Design, www.doschdesign.com). All resulting images were gray-scaled and had a resolution of 256×256 pixels.

The training image dataset consisted of 100 images per object (800 images in total), whereas the test image dataset consisted of 80 images per object (640 images in total). Training image parameters were sampled from a wider range than for the test images (see FigS1B for the sampled distributions). Training images were thus more varied than test images in their position, rotational properties and size.

Finally, the canoncial views for each of the 3D object were defined in prior work (Hong et al., 2016) to align with the common views of that semantic category (Fig 1B). In brief, animals were shown facing forward, with their head upright. The car was shown with the car grille forward and its tires on the bottom. The chair has its legs facing downward while the seat is facing forward. The person is looking straight at the viewer with the top of the head oriented upward. The apple was shown with its stem on top, from a arbitrarily picked rotation (due to its rotational symmetry around the vertical axis). The plane’s cockpit faced, in an upright position.

### Monkey training and behavioral tasks

#### Fixation training and the passive viewing task

We trained all monkeys to maintain fixation on a central fixation dot using operant conditioning with liquid rewards. Specifically, monkeys received rewards for fixating within a square fixation window (2 or 2.5°). We monitored eye movements using video-based eye tracking (SR Research EyeLink II and 1000).

Monkeys initiated a trial of the passive fixation task by looking at a fixation dot. After initiating fixation (300 ms), we presented a sequence of 5-10 object images, each shown for 100 ms at the center of gaze (8°) and followed by a gray blank screen of 100 ms (see Fig 2A for a schematic). If the subject maintained fixation inside the trained fixation window for the entire duration of the image sequence, the image sequence was followed by a fluid reward and an inter-trial interval of 500 ms. All neural data was collected during this passive viewing task to create similar viewing conditions for both groups.

#### Category training and active binary object discrimination task

The category-trained group was additionally trained to perform a binary (match-to-sample) object discrimination task in their home cages using a touch screen (Fig 1A). The full procedure is described in (Rajalingham et al., 2015, 2018) and details on the code and hardware are available at https://github.com/dicarlolab/mkturk and (Ramezanpour et al., 2024). In brief, subjects initiated a trial by touching a central white circle centered on the bottom third of the screen. This triggered the presentation of a sample image for 100 ms presented at the center of the screen (subtending approx. 6-8°). A choice screen immediately followed the sample image. The choice screen contained two object images in their canonical view (the same canonical images were presented across all trials, Fig 1B). One of these objects (Target) matched the category of the object in the sample image. The other (Distractor) was chosen randomly from the seven remaining objects. Monkeys were allowed to respond immediately by touching anywhere within the pixel boundaries of the choice object images. Selecting the correct image resulted in the delivery of a fluid reward, whereas an incorrect choice resulted in a 3 s timeout. Monkeys transitioned to the testing phase once their training performance saturated (for details on this criterion, see Rajalingham et al., 2015). The training phase contained 100 rendered images per object (N=800 images).

During the testing phase (i.e., *active binary object discrimination task*), monkeys saw novel images of the trained categories (N=640 images, 80 per category) in a head-fixed setup using their fixations to make their choices instead of their hands. Specifically, behavior was tested while monkeys viewed images on a 24-inch LCD monitor (1920 x 1080 at 60 Hz, positioned 42.5 cm in front of the animal). To initiate a trial, monkeys fixated a white dot (0.2°) for 300 ms. A sample image was then presented centrally for 100 ms (8°), followed by a blank screen for 100 ms and the choice screen. During the choice period, monkeys could freely move their eyes (for up to 1500 ms). Monkeys made a choice by fixating on the selected image for at least 400 ms. If the monkeys selected the target object, they were rewarded with a fluid reward. A trial was aborted if a subject broke fixation anytime before the choice screen (i.e. if a subject’s gaze was further away than 2° from the central fixation dot).

### Large-scale multi-electrode recordings in the inferior temporal cortex

#### Surgical implants and chronic micro-electrode arrays

All monkeys were surgically implanted with a head-post under aseptic conditions and with up to three 10×10 microelectrode arrays (Utah arrays, Blackrock Microsystems) per hemisphere in IT in a separate surgery. Each array contained 96 electrodes (excluding corner electrodes), each 1.5 mm long and with a 400 μm spacing between electrodes. The placement of arrays was guided by the visible sulcus pattern during surgery (see Fig S2 for the locations per subject and hemisphere). For subjects with recordings from both hemispheres, arrays were first implanted and recorded in one hemisphere, and after approximately a year, these arrays were explanted, and new arrays were implanted in the other hemisphere.

#### Electrophysiological recordings

All neural activity was filtered using an online band-pass filter. Most data contained multi-unit activity and multi-unit spikes which were detected after data collection. In particular, a spike event (multi-unit) was detected when the voltage (falling edge) exceeded more than three standard deviations of the filtered raw voltage values. Firing rates in response to a specific image were obtained by averaging the peristimulus time histogram spike counts (10 ms bins) between 70 and 170 ms after image onset.

In both groups, arrays sampled a variety of regions along the posterior-to-anterior IT axis (Fig S2). Rather than focusing on potential anatomical differences, we here focused on the inferior-temporal cortex as a whole, treating each site as a random sample from the overall IT population. This choice was informed both by prior studies linking IT responses to object recognition behaviors (e.g., Majaj et al., 2015), as well by statistical considerations. In particular, we reasoned that combining all measurements across IT would give us the highest signal-to-noise ratio, an aspect we deemed particularly important given the reported variability of effect sizes in the literature (e.g., Op de Beeck & Baker, 2010).

During recording sessions, we monitored the monkeys’ eye movements using video-based eye tracking. At the start of a session, monkeys performed an eye-tracking calibration task by making a saccade to spatial targets and maintaining fixation for 500-700 ms. We repeated this procedure if a drift in the eye-tracking signal was noticed during a session.

### Neural and behavioral data analyses

#### Behavioral metrics

We quantify behavioral performance as the image-level discriminability reflecting how well a given category can be distinguished from all other possible categories in a given image (similar to (Kar et al., 2019; Kar & DiCarlo, 2021) and defined in (Rajalingham et al., 2018). Specifically, discriminability for an image *i* is defined as:

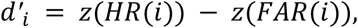

where the hit rate (*HR*) is the proportion of trials for which an image *i* was correctly classified as category *c*. The false alarm rate (*FAR*) for category *c* refers to the average proportion of trials in which any image of another category was incorrectly assigned to category *c*. Finally, *z* refers to the inverse of the cumulative Gaussian distribution.

To estimate the reliability of this image-level metric for the monkey’s behavioral data, we compute the Spearman-Brown-corrected split-half reliability across 20 repetitions for random splits of trial repetitions per image.

#### Neural recording quality metrics and pooling

We aimed to assess group-level differences between task-trained and -naïve recordings at the population level. To ensure that potential differences did not stem from other factors, such as a difference in the number of sites in every group or reliability, we analyzed the recording sites in pseudo-random pools drawn as bootstraps from all recording sites (within a given group). For every draw, we first resampled all recording sites (with replacement) for all subjects. Then, we constructed a single pool using these resampled sites for every group (naïve or trained). To construct the pools, we sampled an equal number of sites per subject (without replacement) as well as sites with an evoked split-half reliability response of at least 0.3 (mean across 1000 repetitions, spearman-brown corrected). This procedure could still produce group differences in reliability since the reliability distributions differ across datasets (Fig S3A). To create IT pools comparable in reliability, we, therefore, sampled recording sites using a weighting based on the untrained reliability distribution (estimated using a Gaussian kernel density estimate; see Fig S3B). We repeated this sampling procedure 1000 times, resulting in 1000 neuron pools for each group with comparable reliability (median: -0.002, see Fig S3C). Within these pools, neural changes were quantified on the repetition- and trial-averaged firing rates of the first 36 image repetitions.

#### Site-level category selectivity

To investigate whether the examined neural recordings differed in how strongly any site modulated its response to the trained categories, we computed a category selectivity index for every recording site (similar to Freedman et al., 2003). For a given site *u* in a pool, we computed the pairwise absolute differences for every pair of images. Next, we averaged the absolute pairwise distances between images from the same object (<u>*SO_u_*</u>) and those of different objects (<u>*DO_u_*</u>). To obtain an index of category selectivity *S_u_*, we finally compared the difference between those two averages and normalized this difference by their sum:

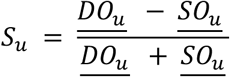

To compare task-trained and -naïve distributions, we summarized all selectivity indices using the median within a pool, thus providing 1000 estimates of selectivity in every group.

#### Population-level categorical representational similarity

We adopted representational similarity analysis (RSA, Kriegeskorte et al., 2008) to assess whether the relative representation of categories is altered by category training. Computing the neural representational distances and comparing them to an ideal categorical model provides a useful additional perspective to linear decoding techniques since it enables us to inspect whether images from the same category show an increased tendency to cluster beyond their overall separability (from other categories).

We first computed the pairwise distances of responses across all sites and images using the Euclidean distance to estimate categorical representational similarity. This yielded a symmetric representational dissimilarity matrix across images for every pool. In a second step, we computed the second-order correlation of the lower triangle (excluding the diagonal) with a categorical RDM (assigning full dissimilarity to images of different categories and zero dissimilarity to those of the same category). Since the categorical RDM is ordinal, we adopted Kendall’s tau for the second step with a correction for rank ties to avoid inflated estimates (Schütt et al., 2023).

#### Population-level category decoding

To estimate how much information can be linearly decoded from IT neural populations by downstream neurons receiving inputs from IT, we used linear classifiers, which have previously been shown to sufficiently link IT population responses to human behavior (Majaj et al., 2015).

We optimized linear classifiers (one-vs-all) on the image-evoked time-averaged rates in a given pool to predict image categories of the shown images. Specifically, we fit logistic regression classifiers with L2-regularization using a 3-fold cross-validation procedure with stratified random splits (across categories). This procedure provided held-out category predictions for all images. The inputs to the regression models were scaled based on the training data split. We repeated this procedure 20 times (for different data splits) and computed image-level *d’* based on all predictions (see *Behavioral metrics).* We adopted the mean performance (image-level *d’*) across repetitions in further analyses. Regression models were implemented in scikit-learn (Pedregosa, 2011) adopting default parameters.

#### Population-level decoding of category-orthogonal properties

In addition to category-related information, we assessed group-level differences in the available information about category-orthogonal properties (object location, eccentricity, size, and rotation, similar to Hong et al., 2016). Note that the properties were also used during the image generation process, and therefore, ground-truth values were available for all properties.

To estimate information about category-orthogonal properties, we followed the same approach as for category decoding with two key differences: To accommodate the prediction of continuous properties, we used Ridge Regression (implemented in scikit-learn (Pedregosa, 2011) and adopting default parameters) and performance was quantified as the Pearson correlation between the held-out predictions (cross-validation) and the ground-truth image properties.

#### IT-to-Behavior consistency

We probed the alignment between monkey image-level performance (behavior, *b*) and IT decodes (neural, *n*) by estimating the image-level consistency between behavior and neural responses. Specifically, we computed the Pearson correlation between the image-level behavioral and IT category *d’* and then normalized this correlation by the geometric mean of their spearman-brown-corrected split-half reliabilities:

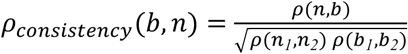

We repeated this procedure 20 times with different split halves of trials (i.e., repetitions of an image) to estimate the reliability of this measure.

#### Simulation of monkey training using artificial neural networks

We built a variety of simulations of the monkey’s training to ask whether any changes observed empirically between trained and naïve IT responses are consistent with updates to current computational models of the visual ventral stream.

#### Base models

As our base models of task-naïve monkey IT and behavior, we considered image-computable artificial neural networks optimized on large-scale image databases previously shown to be predictive of IT responses and categorization behavior (e.g., Brain-Score platform, Schrimpf et al., 2018). To reflect that the pre-training of these models might influence their fit to the empirical data, we included models optimized with supervised training on ImageNet (ResNet18, 34, 50, 101, 152, VGG16, 19, and AlexNet), weakly-supervised pre-training followed by supervised training on ImageNet (resnext101_32x8d_wsl, resnext101_32x16d_wsl) and self-supervised training on ImageNet (ResNet50 trained with MoCov2, barlowTwins and SimCLR). All supervised and weakly-supervised base models were retrieved from Pytorch (Paszke et al., 2019), whereas self-supervised models were obtained from VISSL (Goyal et al., 2021).

#### Aligning base models to category-naïve IT responses

To ensure that potential changes observed between task-naïve and -trained IT responses are comparable in scale and feature complexity in the considered base models, we first aligned the candidate models (before training) with the untrained IT sites. To achieve this, we identified the layer in all considered base models with the highest encoding capabilities of IT responses using standard procedures in the brain-score framework. Specifically, we computed a cross-validated PLS regression between a layer’s activation features and IT responses, providing a predictivity score for all evaluated layers. In all analyses, we simply picked a base model’s layer with the highest score as our IT-mapped layer (see Fig S5 for an base model overview, incl. the layer scores).

It is an open question how a single unit in an ANN model layer relates to a single recording site in a monkey’s IT cortex. Previous work on linking ANNs to neural responses has adopted various approaches (e.g., adopting all layer units and performing dimensionality reduction, e.g., Conwell et al., 2023). Here, we attempted to mimic the sampling process of the neural recordings as closely as possible rather than assuming access to the information of all layer units. To this end, we assessed at what scale (units per recording site) the decodable information from randomly drawn units in a base model’s IT-mapped layer is comparable to the average decoding accuracy observed in task-naïve IT recording sites (on average across 300 pools, 159 sites per pool).

To achieve this, we first sampled category decodes (see *Population decode accuracies*) for different scaling factors in the IT-mapped layer using a log-scale sampling (1-100 unit scaling, equivalent to 159-15900 randomly sampled units). In a second step, we then fit these data points using an x-shifted log function, where *a*, *b*, and *c* were free parameters:

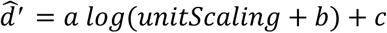

Fitting this function to model simulations using different scaling factors gave us a continuous function between the number of randomly sampled units in the IT-mapped layer and its decodable information (*d’*) (see Fig S6A). Finally, using this fitted function per IT-mapped base model layer, we identified the scaling factor equivalent to the task-naïve IT recording decodes (Fig S6B). If a base model’s IT-mapped layer failed to reach this criterion within the sampled range, we excluded that base model from further analysis. We found that none of the weakly-supervised IT-mapped layers (resnext101-32x8d-wsl and -32x16d-wsl) reached performance levels on par with task-naïve IT decodes. As a result, we excluded these base models from the training simulations.

#### Training strategies

How monkey training compares to training an artificial neural network is an open scientific question that touches on many current issues, including what mechanisms support different forms of plasticity in the brain or whether back-propagation is a proxy of biological learning. In this work, we sought to make this comparison tractable by focusing our efforts on currently available training approaches capable of yielding performance improvements in object classification for the training images seen by the trained monkeys.

We matched the monkey and model training parameters wherever possible and included a wide range of parameters when the analogy was less clear. To mimic the monkey’s training, we only included images the monkeys have seen during training (N=800). Furthermore, we optimized any models for 15 epochs, equivalent to an exposure of 1500 repetitions per category, previously shown to be a typical timescale for monkeys to learn new categories (Rajalingham et al., 2015). For any choices where we could not draw an analogy to the monkey training, such as the learning rate, batch size, regularization, and the optimizer’s momentum, we set up a search space with parameter ranges informed by common choices in machine learning, reasoning that this would produce a variety of performant models with diverse parameter configurations. We adopted a Bayesian hyperparameter search implemented via the weights and biases framework for 20 repetitions (Biewald, 2020).

Finally, we adopted a few standard practices for optimizing neural networks: we optimized all models using a cross-entropy loss function, a stochastic gradient descent optimizer, and a step decay learning rate scheduler (reducing the learning rate every six epochs by a factor of 0.1). We dedicated 30% of the training images as a validation set. This enabled us to choose the best model across training epochs for further evaluation. Beyond these choices (common to all trained models), we devised three training strategies:

##### Standard finetuning

We replaced the output layer of a pre-trained model with a randomly initialized fully connected layer. During training, we provided 1-hot feedback across all eight training classes. The resulting error gradients could modify any weight in the pre-trained network.

##### Step-wise finetuning

This approach was identical to standard finetuning, with the only difference being that we initialized the fully connected layer with an optimized linear regression readout. This approach yields higher-performing fine-tuning results and improved generalization by circumventing large errors early in training (Kumar et al., 2022). This training choice was motivated by the observation that common category training paradigms typically span multiple days and thus may feature both within and between-session improvements (Op de Beeck & Baker, 2010).

##### Binary choice

This strategy mimicked the feedback structure in a delayed match-to-sample task with two choices used for monkey training. In contrast to the feedback provided during supervised training specifying both the target as well as the absence of all distractors (1-hot encoding, information on all seven distractors), monkeys only received feedback on one target and one distractor class in a given trial. To simulate this form of sparser feedback, we modified the gradients such that a model only received feedback on the target and one randomly chosen distractor class for a given image.

All considered candidate models were optimized 20 times for every training strategy, resulting in 660 trained models (11 models, 3 strategies, 20 repetitions).

#### Model selection

We selected trained models for a detailed comparison with the empirical data if the trained model reached a behavioral performance level (average *d’* across images) on par with the trained monkeys (i.e., within the 95% confidence interval of the hierarchically pooled monkey behavior).

#### Quantifying changes in candidate models

To assess the changes in the IT-mapped layers, we adopted the same three category metrics used during neural data analysis (i.e., category selectivity, linear decoding, and categorical representational similarity). All metrics were computed for 100 randomly drawn unit pools. For site-level category selectivity, we adopted the average response of drawn units (at the specified site-to-unit scaling) as a proxy of site-level selectivity.

#### Comparing model and neural differences

To effectively compare neural differences to those of the candidate models, we first computed the relative difference for all metrics (*ϕ*) in both models and the neural data:

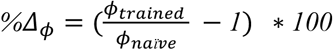

This estimates the difference (*Δ%*) for every pair of draws (from the task-trained and -naïve pools, respectively) for the neural data and the IT-mapped layers. We averaged across the repeated draws (from the IT-mapped layer) to obtain a model’s *Δ%* estimate. We considered models as IT-like if the *Δ%* estimate across all metrics fell within the 95% confidence interval of the observed changes in the IT data (Fig 5C). In turn, models that only matched a subset of metrics were less IT-like, and models that matched none of the metrics were referred to as IT-unlike.

#### Statistical analyses

To test for differences between task-trained and -naïve IT pools, we adopt two increasingly conservative approaches: First, to test for differences between groups (1000 draws sampled pools with replacement), we performed a two-sample bootstrap test by computing the differences in the bootstrapped distributions and derive a *p*-value from the probability of zero under that difference distribution (i.e., we construct the confidence interval of the difference distribution and check whether it contains zero). Second, to account for different sources of within and between-subject variability during our neural pooling procedure (Fig S4), we also performed a hierarchical permutation test comparing differences observed in the empirical data with those expected from pools sampled with permuted group assignment (similar to a hierarchical randomization approach by (Kulkarni et al., 2022). To implement this, we created all possible combinations of groups (while ignoring the experimental conditions) and performed our standard neuron pooling procedure for these groups. This gives a null distribution of differences for the randomly created groups, which we then adopt to interpret our empirically observed differences (i.e., the difference between task-naïve vs. trained pools). Due to the computational demand of this second approach, we reduced the number of draws for any group to 200.

To compare model differences to those in the empirical data, we tested whether the average estimated differences of a model fell within the 95% confidence interval of the empirical data (across draws/bootstraps). This approach was applied to both the behavioral and IT-based metrics. We adopted an alpha level of 0.05 to evaluate statistical significance throughout the project.

## Acknowledgments

This work was funded in part by the National Science Foundation-STC [CCF-1231216 & 2124136] (JJD), Simons Foundation [AN-NC-GB-Culmination-00002986-04] (JJD), Office of Naval Research [N00014-21-1-2801 & N00014-20-1-2589] (JJD) and a DFG Walter-Benjamin Fellowship [547591872] (LKAS). KK has been supported by funds from the Canada Research Chair Program [CRC-2021-00326], the Simons Foundation Autism Research Initiative [SFARI, 967073], the Canada First Research Excellence Funds [VISTA Program], and NSERC Discovery Grant. We thank the members of the DiCarlo lab and, in particular, M.J. Lee, for insightful discussions and advice.

## Supplementary figures

**Figure S1.**
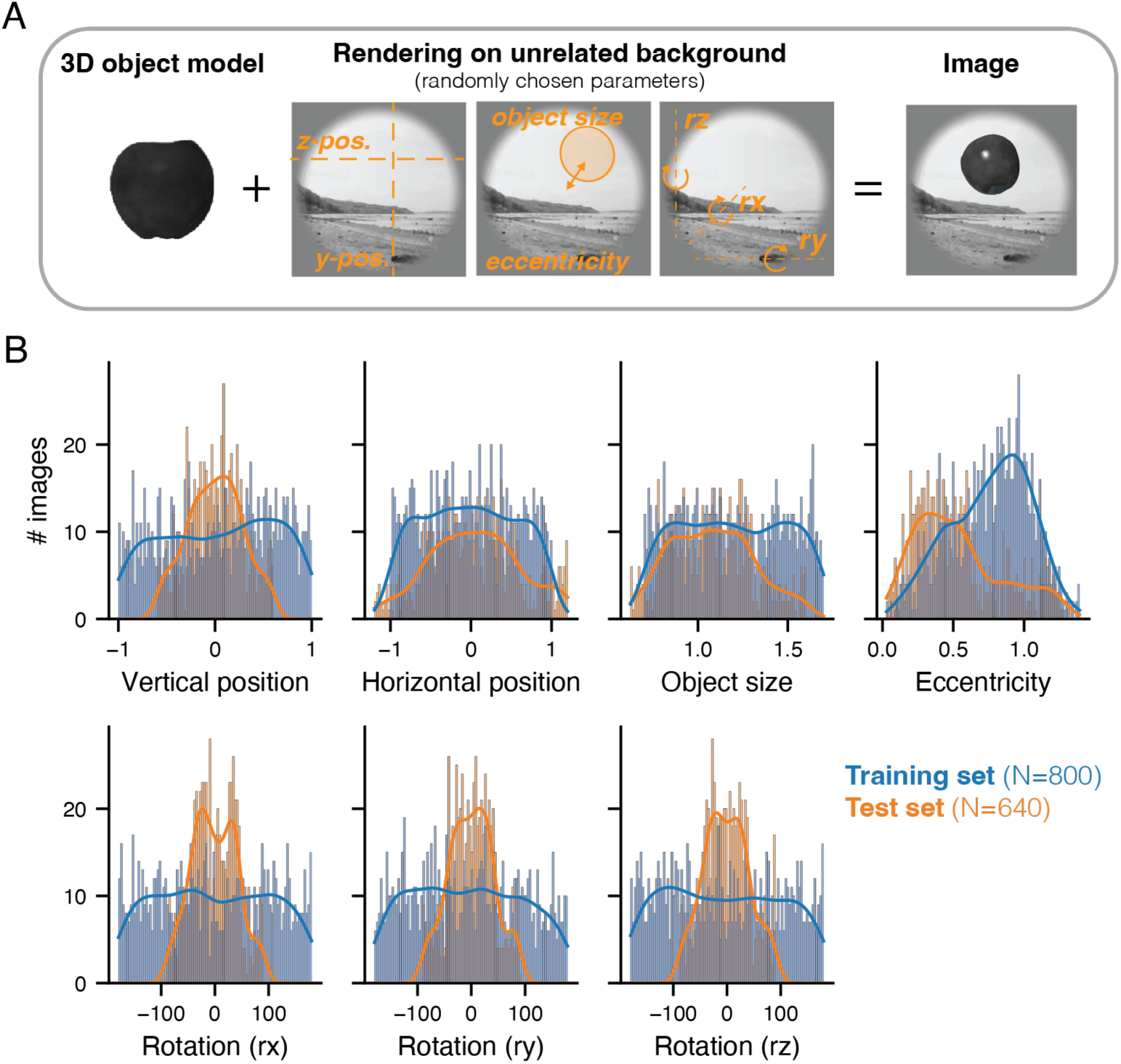
Image dataset generation. (A) Training and test image dataset were generated by rendering a 3D object model on a randomly chosen scene background. The 3D object model was rendered using a randomly chosen set of parameters (within a specified range), yielding a novel object image. (B) Comparison of the training and test image sets across the rendering parameters. Test images were sampled from a narrower parameter range compared to training images. Every image (incl. training and test set) was generated from a unique combination of parameters and there was no link between an object’s identity and its rendering parameters.

**Figure S2.**
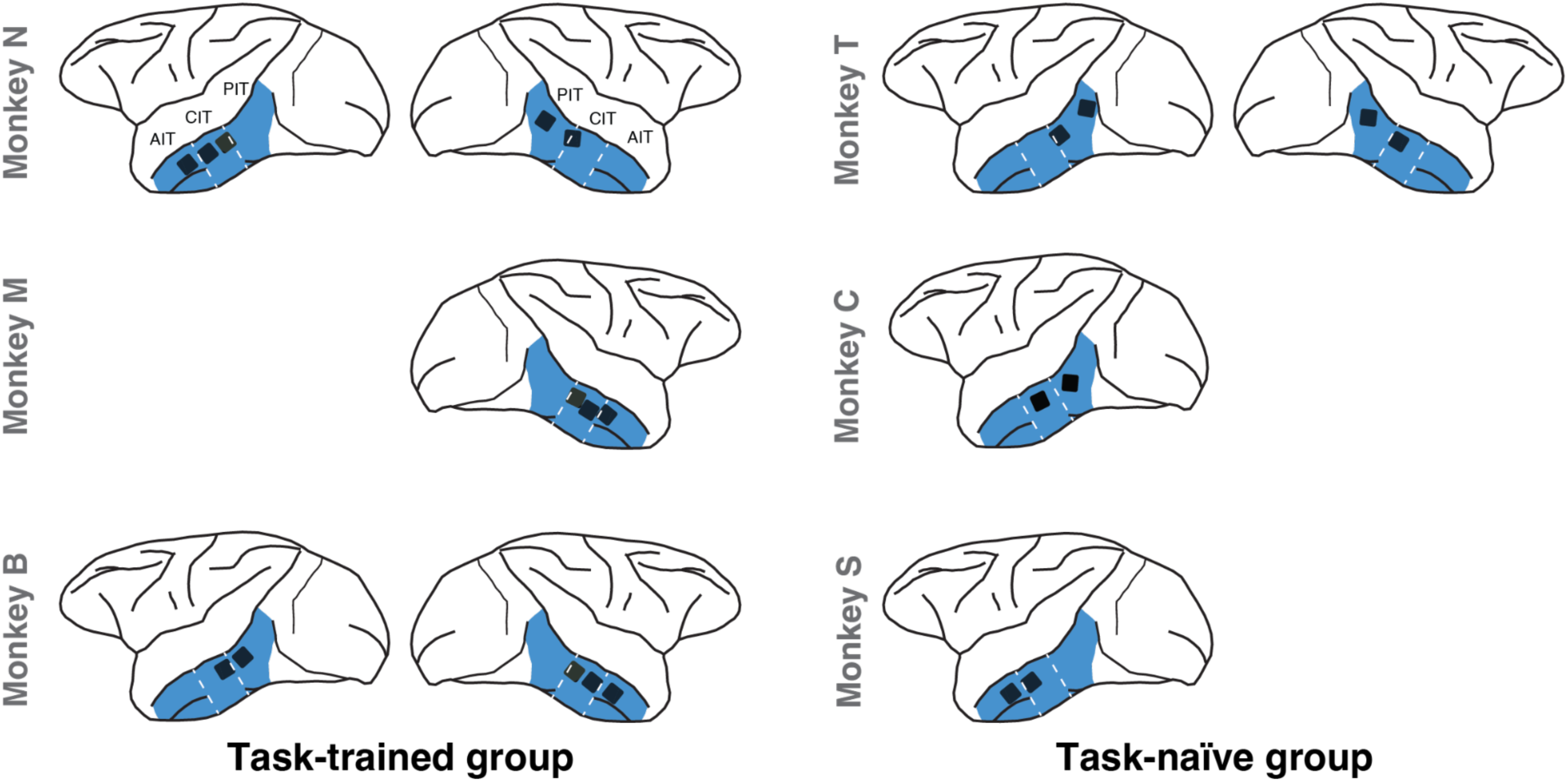
Micro-electrode array placements. All subjects had chronically implanted Utah arrays in various locations along the posterior-to-anterior axis in the inferior temporal cortex. Note that if arrays in both hemispheres are shown, recordings were performed consecutively in both hemispheres (see Methods for further details). Arrays were classified as AIT, CIT or PIT according to their position relative to the anterior and posterior middle temporal sulci (annotated by the grey dashed lines). For our main analysis, we did not take the spatial location of the arrays into account and treated all recording sites as random samples from an average IT population.

**Figure S3.**
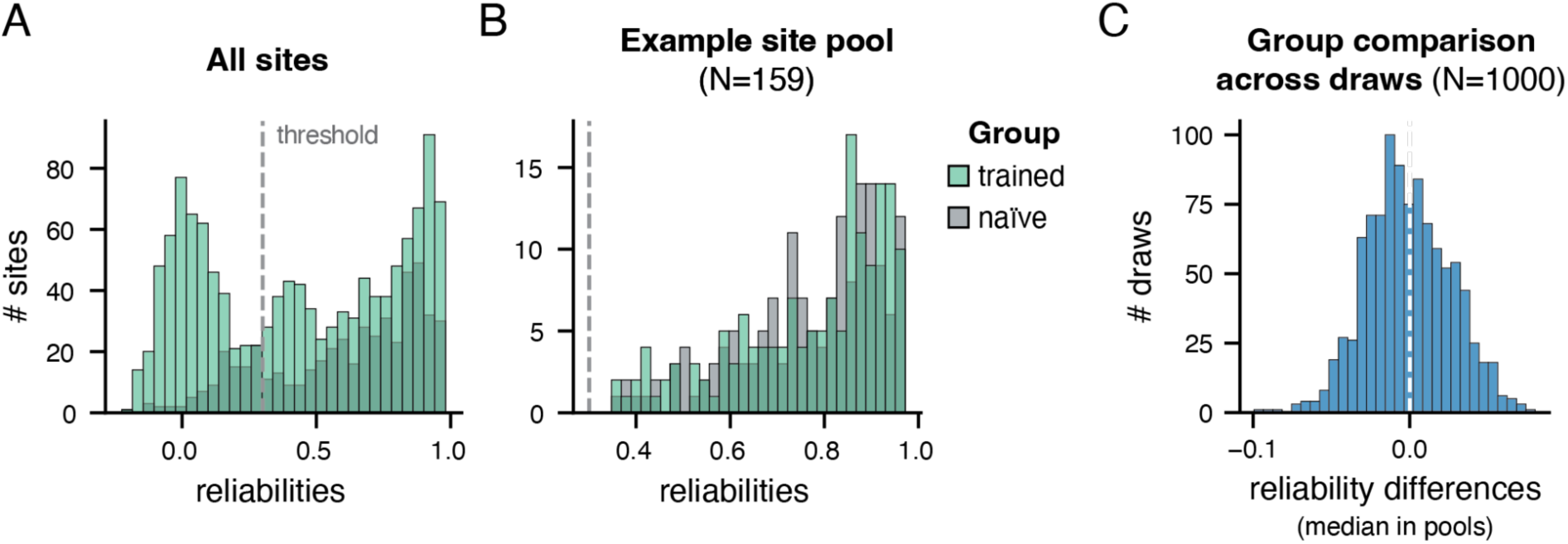
Matching site reliability across groups. (A) The IT recordings collected from task-trained and -naïve subjects differed in the number of available sites and the reliability distribution across sites. We computed reliability as the correlation across averages from random split-halves of trial repetitions (N=1000). The dashed line shows the applied reliability threshold of 0.3. (B) Our pseudo-random pooling procedure (see Methods) produces pools of recording sites corrected for reliability and number of sites per subject (53 sites/subject, thus 159 sites/pool). (C) Using this procedure, we obtain pools that are comparable in their reliability. A total of 1000 pseudo-random pools are shown resulting in a median of -0.002 across all draws.

**Figure S4.**
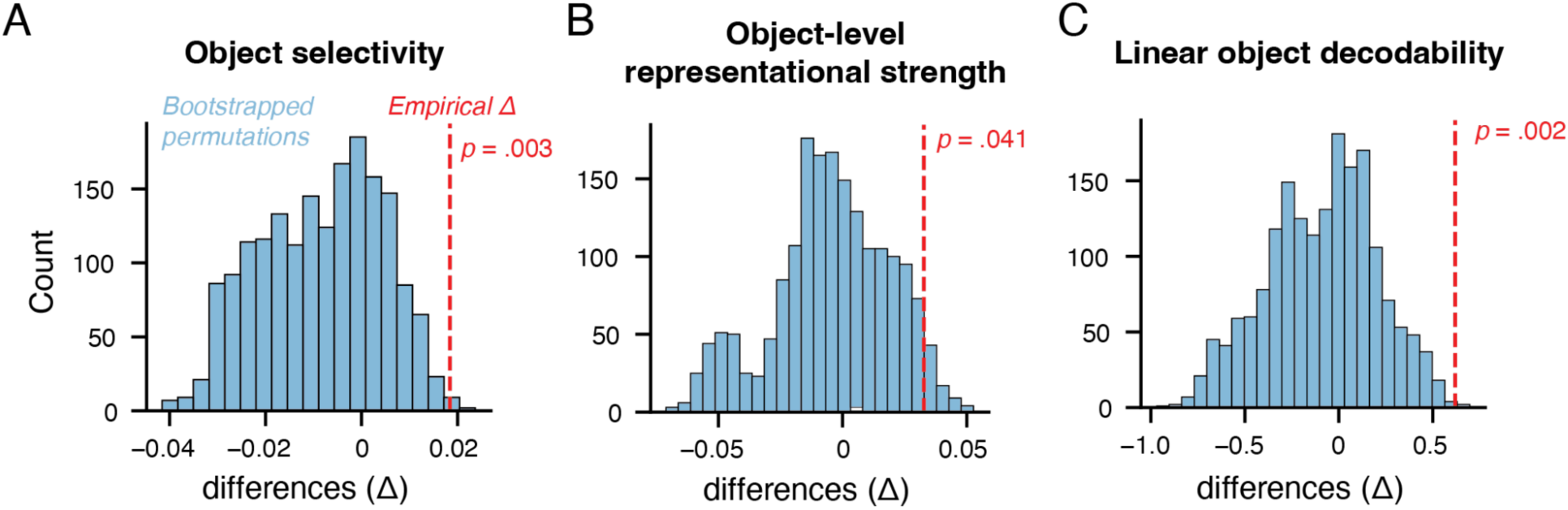
Hierarchical permutation bootstrap test. To what extent are the observed differences driven by differences between a few subjects irrespective of their training history? To control for the effects introduced by interindividual variation in both groups, we sampled a bootstrapped null distribution (200 draws) while permuting the group membership of all monkeys (shown in blue) for all metrics (A-C). Relating the observed differences (empirical, red dashed line) to this null distribution revealed that all observed effects were larger than those expected from interindividual variation.

**Figure S5.**
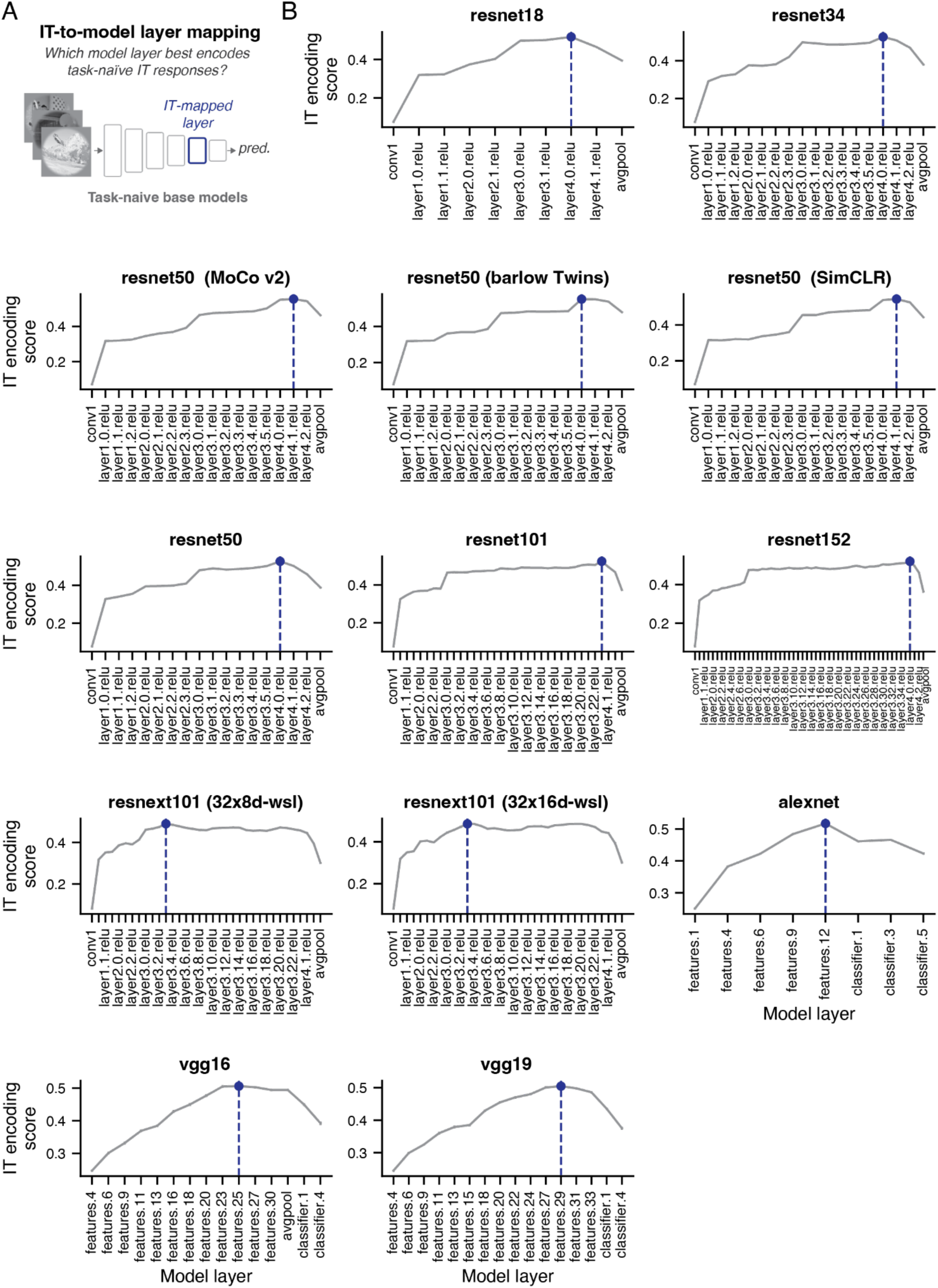
IT-layer mapping for base models. (A) We selected the layer for comparison with the IT data by asking which layer in a given base model best encodes the responses in naïve IT recordings. (B) We evaluated this by adopting the approach taken by the Brain-Score platform. In particular, for any given layer, we fit a cross-validated partial least sqaures regression model to predict neural IT responses. We repeated this approach ten times, each yielding an IT encoding score for a given set of layer features. Here we show the mean across repetitions as a function of layer position within the model (x-axis). The final chosen IT-layer is annotated by the vertical dashed blue line. For visualization purposes, we reduced the number of annotated x-axis labels for all resnet 101 and 152 models.

**Figure S6.**
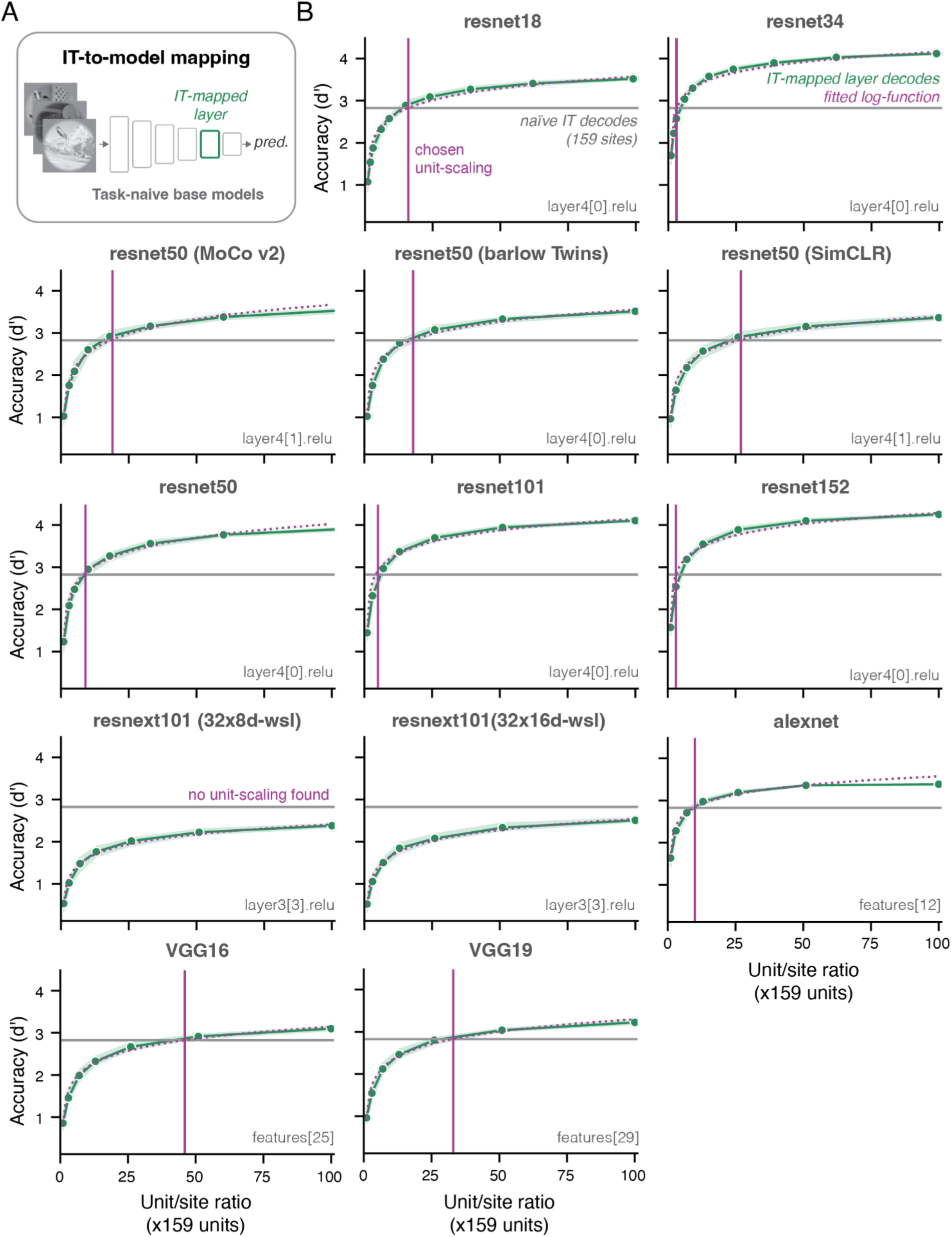
Site-unit scaling for base models. (A) We identified an IT-mapped layer for every base model using the brain-score mapping approach. The chosen layers are annotated in the lower right for all the base models depicted in B. (B) Decoding performance increases as a function of the number of randomly sampled units in an IT-mapped layer for all base models (one subplot each). Performance was estimated using 20 random draws and the error bars show the 95% confidence interval across draws. The x-axis refers to the unit-to-recording site ratio. A ratio of 1 means that 159 units have been randomly sampled. The gray line shows the average decoding performance obtained from task-naïve IT pools (cf. Fig 2K). We fit the neural decodes with a log-function (shown as a dotted line in purple) and then find the intersection of that curve with the untrained performance of IT responses. We adopt the unit-site ratio at this intersection (purple vertical line) as the chosen unit-site scaling for a given base model.

**Figure S7.**
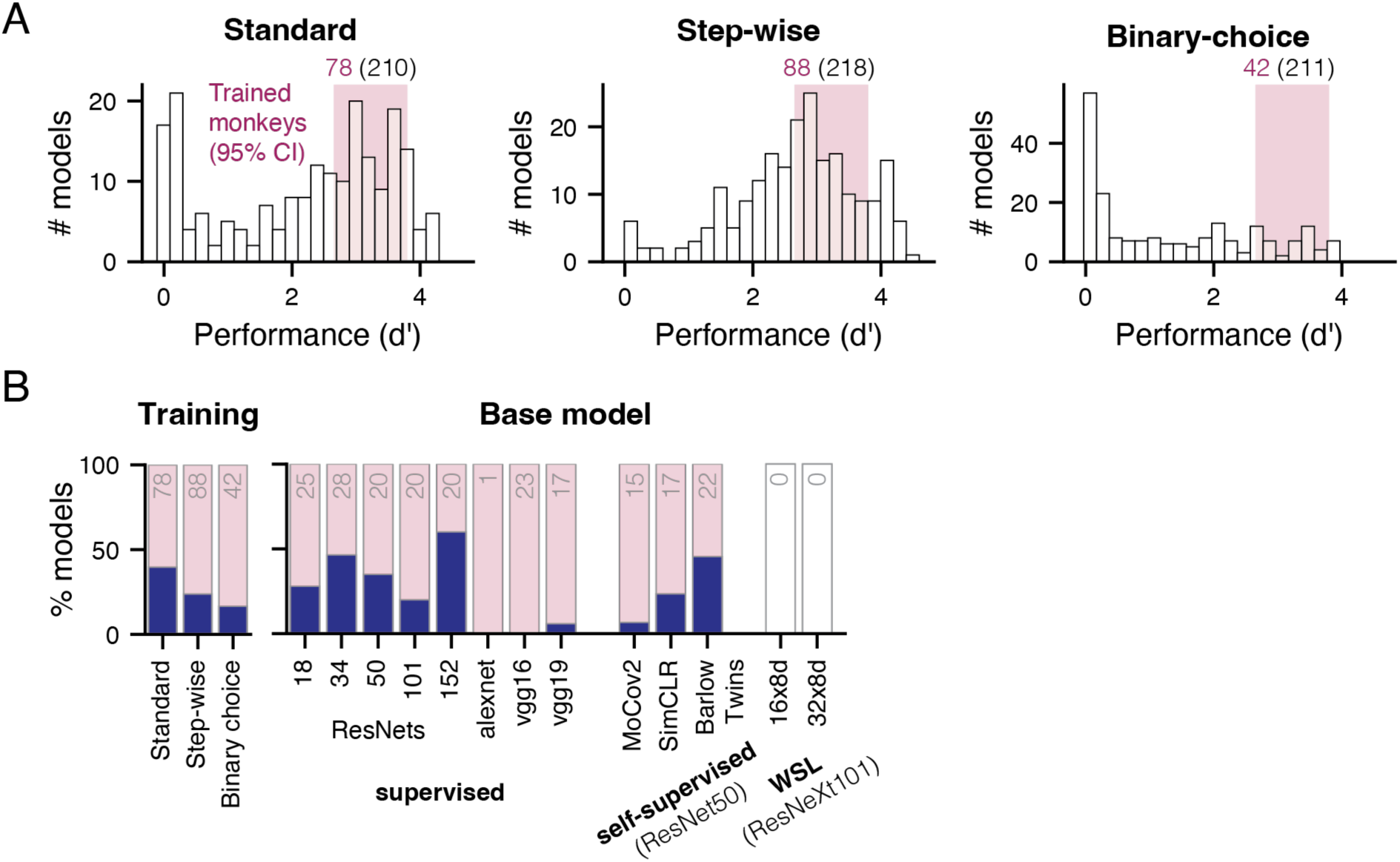
Model inclusion as a function of training parameters. (A) We only considered models as hypotheses of IT plasticity which matched the performance of the trained monkeys (magenta shaded area depict the 95% CI of the pooled monkeys’ behavior). Binary choice training produced the fewest equally performant models, followed by standard and stepwise training. Each training consisted of 20 model iterations (with different combinations of hyperparameters) for the 11 base models for which we could establish an IT-layer mapping. A few model trainings failed, thus producing fewer than 220 models. (B) Which training aspects were associated with producing IT-like models? Every training strategy and most base models could generate models that matched IT along all metrics. The annotated number refers to the number of models that passed behavioral screening or sufficient decoding to establish a mapping with IT responses. The filled blue bar only includes models that match all three metrics.

**Figure S8.**
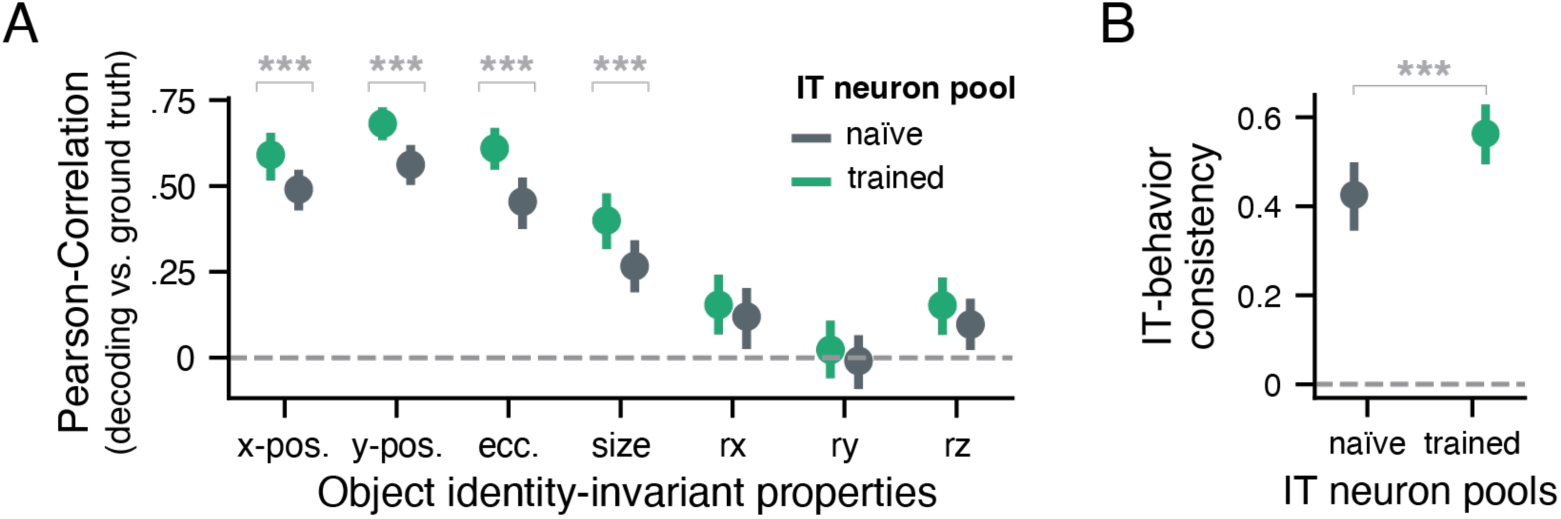
Task training introduces differences in object identity-invariant task decodes and IT-behavior consistency. (A) Decoding accuracy for object identity-invariant image properties (x-axis) from task-naïve and task-trained IT responses. Accuracy is assessed via a correlation between the ground-truth image latents of the 640 test images and the held-out cross-validated decoder predictions (y-axis). (B) IT-behavior consistency between task-trained behavior and task-naïve or -trained IT-object decodes. IT-behavior consistency (y-axis) refers to the noise-corrected correlation between the image-specific performance estimates. In both panels, error bars depict the 95% confidence interval across 1000 pseudo-random draws from the respective neuron pools. All starred comparisons are statistically significant at p < .001.

**Figure S9.**
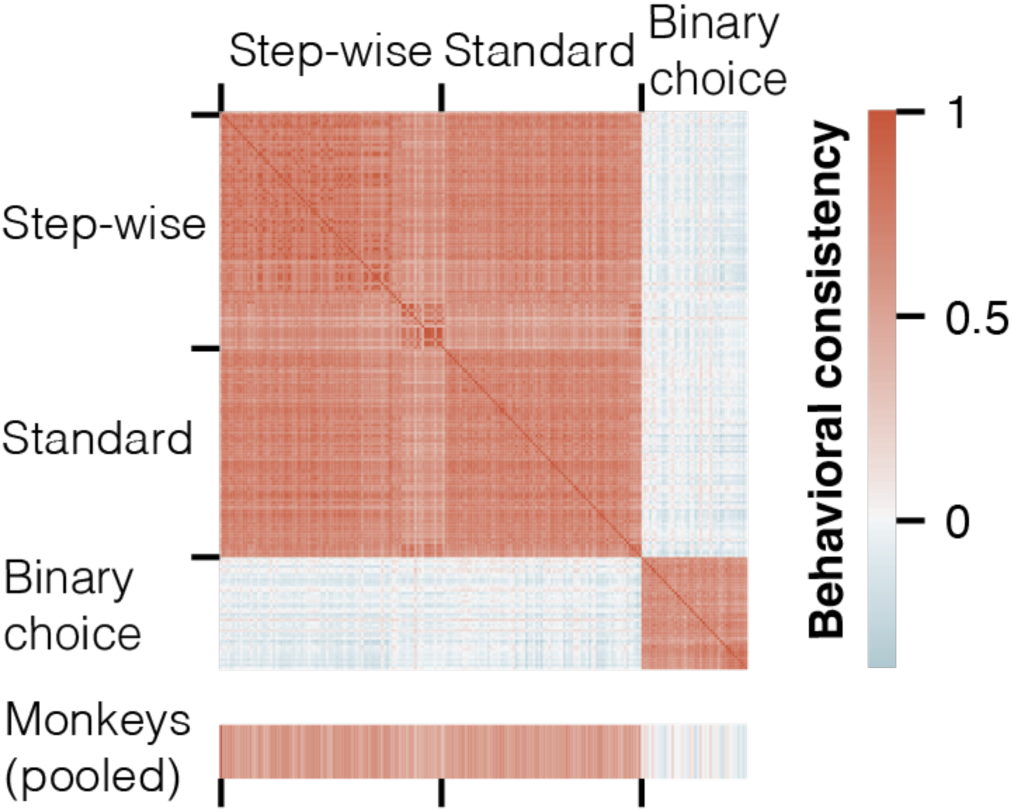
Comparison of image-specific behavioral patterns between trained models and monkeys. Noise-corrected correlations (i.e., consistency) between the image-level behavioral performance across 640 test images for all 208 models and the pooled monkey behavior. Individual models are ordered according to their training approach. While models trained with the step-wise and standard training approach are highly consistent with each other after training, binary choice trained models were only consistent with themselves but not with models from other training approaches. Image-wise performance patterns estimated from the behaviors of 3 trained monkeys are consistent with step-wise and standard training but not with models trained with binary choice feedback. The height of the monkey data is increased to enhance visibility.

## Notes

**Conflict of interest** The author declares no competing financial interests.

### Competing Interest Statement

The authors have declared no competing interest.

### Summary of Updates

This version includes additional analyses (e.g., Fig 3).

